# The importance of the location of the N-terminus in successful protein folding *in vivo* and *in vitro*

**DOI:** 10.1101/2023.12.11.571183

**Authors:** Natalie R Dall, Carolina A T F Mendonça, Héctor L Torres Vera, Susan Marqusee

## Abstract

Protein folding in the cell often begins during translation. Many proteins fold more efficiently co-translationally than when refolding from a denatured state. Changing the vectorial synthesis of the polypeptide chain through circular permutation could impact functional, soluble protein expression and interactions with cellular proteostasis factors. Here, we measure the solubility and function of every possible circular permutant (CP) of HaloTag in *E. coli* cell lysate using a gel-based assay, and in living *E. coli* cells via FACS-seq. We find that 78% of HaloTag CPs retain protein function, though a subset of these proteins are also highly aggregation-prone. We examine the function of each CP in *E. coli* cells lacking the co-translational chaperone trigger factor and the intracellular protease Lon, and find no significant changes in function as a result of modifying the cellular proteostasis network. Finally, we biophysically characterize two topologically-interesting CPs *in vitro* via circular dichroism and hydrogen-deuterium exchange coupled with mass spectrometry to reveal changes in global stability and folding kinetics with circular permutation. For CP33, we identify a change in the refolding intermediate as compared to WT HaloTag. Finally, we show that the strongest predictor of aggregation-prone expression in cells is the introduction of termini within the refolding intermediate. These results, in addition to our findings that termini insertion within the conformationally-restrained core is most disruptive to protein function, indicate that successful folding of circular permutants may depend more on changes in folding pathway and termini insertion in flexible regions than on the availability of proteostasis factors.

## Introduction

In the cell, most proteins have their first opportunity to fold during their synthesis via translation. Protein synthesis occurs orders of magnitude slower than the rate of formation of α-helices and β-strands, which allows for sampling of various conformations as the nascent polypeptide chain emerges from the ribosome (1–5). Recent studies of this co-translational process have demonstrated secondary structure formation within the ribosome exit tunnel and during translation (co-translational folding) (6–10). The vectorial nature of protein synthesis from N- to C-terminus generates a potential bias in the conformations available to the growing nascent chain, with the ability to influence the overall folding trajectory and efficiency (11–13). Several proteins have been shown to fold more efficiently (producing more soluble product) co-translationally as compared to refolding *in vitro*. For example, firefly luciferase and GFP are both highly aggregation-prone during refolding, yet efficiently produce soluble protein when translated *in vitro* or when over-expressed in cells (14–16).

*In vivo*, the folding process is also modulated by cellular factors — chaperones and other parts of the proteostasis network (17). For example, soluble refolding of firefly luciferase can be rescued by introducing the bacterial chaperone system DnaK/DnaJ/GrpE to *in vitro* refolding experiments (18). Chaperones such as trigger factor and DnaK/DnaJ (HSP70 and HSP40 in mammalian systems) can interact with nascent chains to block the formation of nonproductive, kinetically-trapped intermediates during co-translational and post-translational folding (19–21). Protein expression can also be modulated by translational speed, where slowing or increasing the rate of translation by introducing and shuffling rare codon positions have been seen to impact expression yields and folding pathways (22–24). Thus, folding in the cell is highly dependent on several factors external to the sequence of the growing polypeptide chain. Despite this, we know very little about the importance of which part of the protein is translated first (that is, where the N-terminus resides within the overall fold of the protein).

The order of protein translation can be easily modified via circular permutation. Circular permutants (CPs) are created, often at the DNA-level, by connecting the N- and C-termini with a flexible amino acid linker and introducing the termini elsewhere in the protein (25–27). For proteins where the natural N- and C-termini are close in the three-dimensional structure, the circular permutant is often structurally-similar or identical to the wild-type protein. CPs also often retain function, even though the vectorial synthesis of the nascent chain during translation is altered. It is important to note that although the native structure may be resilient to permutation, the energy landscape of the protein may be drastically perturbed. Circular permutation has been observed to affect aggregation propensity, quaternary structure, catalytic activity, folding, and even proteolytic susceptibility in different cell types (28–32). Importantly, circular permutants have the potential to form novel co-translational folding intermediates not observed in the wild-type protein, as different regions of the protein will be synthesized first on the ribosome for each CP. Thus, permutation has the potential to bias the folding pathway, with CPs populating or avoiding off-pathway intermediates, affecting misfolding and/or aggregation. This, in turn, may alter protein interactions with chaperones and other members of the proteostasis network, although to date, we are unaware of any experimental documentation of such.

In previous studies, work from our lab demonstrated that co-translational folding of the protein HaloTag is less aggregation-prone than when refolded *in vitro* from chemical denaturant (33). This co-translational folding was monitored in an *in vitro* purified translation system (PURExpress®) in the absence of any cellular factors such as chaperones or proteases. HaloTag is a 297-residue bacterial haloalkane dehalogenase engineered to covalently bind halogenated ligands (34). Similar to other members of the α/β hydrolase protein family, HaloTag contains a core subdomain made up of an α/β sheet with eight β-strands connected by α-helices, and a helical lid subdomain inserted between β6 and β7 (35, 36) (Fig. 1a). When refolded from denaturant, HaloTag populates an aggregation-prone refolding intermediate with structural elements from both N- and C-terminal regions (33). These results suggest that the vectorial nature of translation may bias local structures populated while still a nascent chain on the ribosome and thereby avoid the formation of this aggregation-prone partially folded structure, as the C-terminal intermediate residues will be sequestered in the ribosome until translational termination is complete.

**Figure 1.**
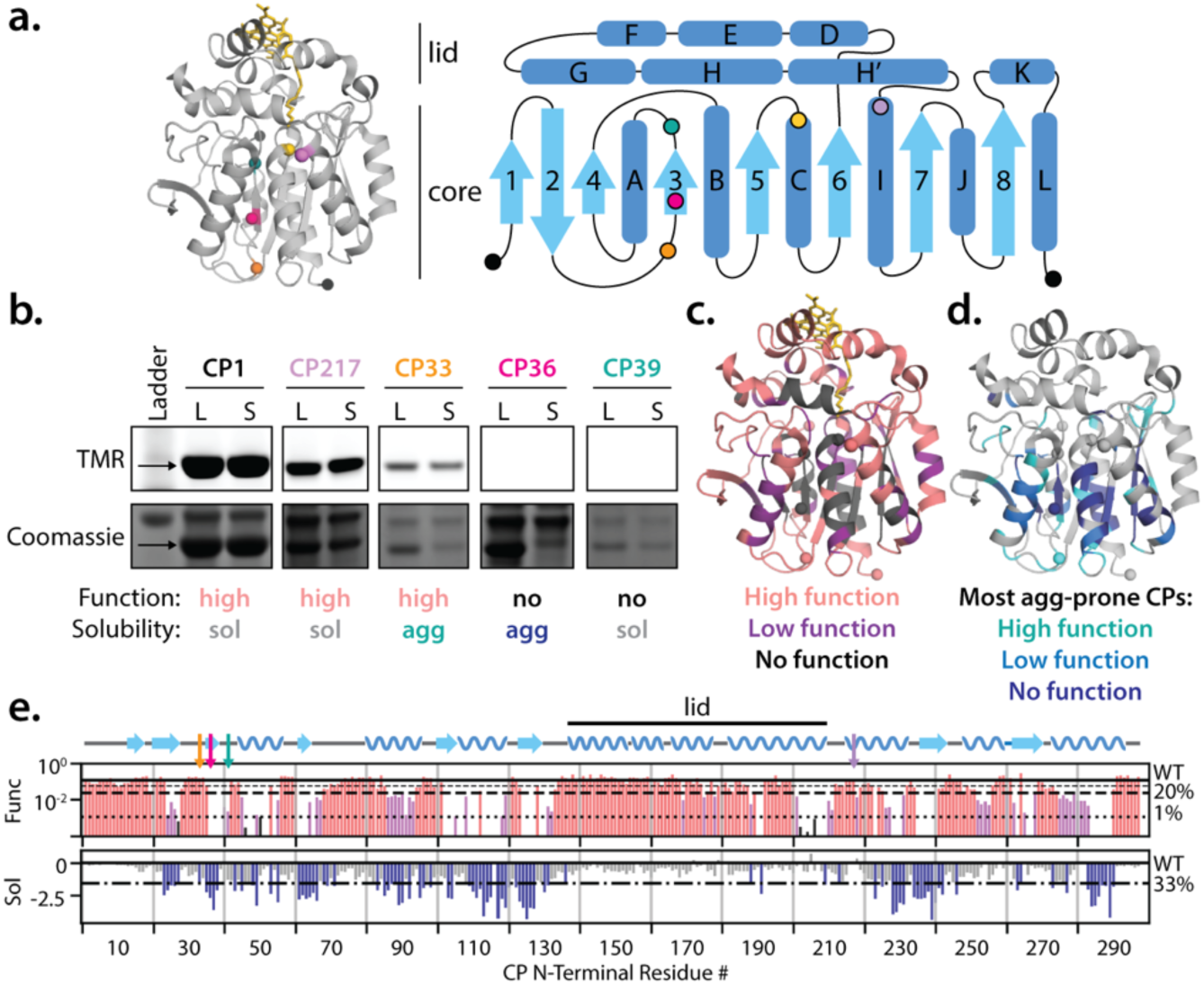
Circular permutation of HaloTag impacts function and solubility in *E. coli*. a. Left: Structure of HaloTag bound to the ligand TMR (yellow sticks, PDB: 6u32). Right: Secondary structure topology map of HaloTag. Helices shown in dark blue, β-strands shown in light blue. Spheres: native termini (black), CP33 (orange), CP36 (dark pink), CP39 (teal), and CP217 (purple). Yellow sphere indicates D106, the catalytic residue covalently-attached to TMR. b. Representative gel-based assay using a detergent-based lysis method examining solubility and function of HaloTag CPs. CPs showing TMR fluorescence in top gel image are considered functional. CPs showing equal amounts of protein in the supernatant fraction (S) as the whole cell lysate (L) are considered soluble, while CPs less-abundant in the supernatant are considered aggregation-prone. Black arrows show CP bands. c. Locations of functional (light pink), slightly-functional (dark pink), and nonfunctional (purple) CPs. d. Locations of the most aggregation-prone CPs. Teal CPs are highly aggregation prone yet functional, while dark blue CPs are aggregation-prone and nonfunctional. e. Quantification scores based on the gel assay for every possible CP. One-dimensional topology map is shown above, with colored arrows indicating positions of CPs highlighted in a-b. Functional scores are calculated based on the relative amounts of TMR:protein based on gels quantified in ImageJ. Lines indicate the WT score(-), 20% of WT (--), and 1% of WT (••). Functioning CPs have scores above the 20% line (pink), low-function CPs have scores between 1-20% the WT score (purple), and nonfunctioning CPs have scores of 0-1% the WT-level (black or no bar). Solubility scores represent the ratio of protein in the supernatant and lysate fractions, are normalized to WT HaloTag, and log_2_-transformed, so scores closer to zero indicate higher solubility. Lines indicate the WT score (-) and 33% of WT (-•-). CPs with scores below the 33% WT line (navy) are considered highly aggregation-prone.

Here, we have taken a comprehensive approach to evaluate the effect of circular permutation on a protein, its ability to fold and function, and its reliance on the translational machinery and cellular factors when expressed in cells. We constructed all 297 possible circular permutants of HaloTag and evaluated their function in multiple cellular environments. To quantify solubility and function, we use both a biochemical gel assay with *E. coli* cell lysate and a high-throughput screen of protein function via fluorescence-activated cell sorting coupled with next-generation sequencing (FACS-seq). To evaluate the role of cellular factors, specifically the co-translational chaperone trigger factor and the intracellular protease Lon, we carried out the FACS-seq experiments using *E. coli* cells lacking these specific factors. Finally, we selected two novel circular permutants for detailed biophysical characterization to evaluate the effects of termini relocation on structure, folding, and stability (the energy landscape). Combining these results allows us to identify trends that correlate with the function and solubility of protein CPs, which carry implications for protein engineering, termini placement in protein design, and the evolution of complex protein structures.

## Results

### Circular permutation of HaloTag differentially impacts protein solubility and function

To evaluate the role of circular permutation in protein folding, we generated a library of 297 plasmids encoding all potential circular permutants of the protein HaloTag (Supp. Fig. 1a). For each permutant, an eight-residue linker (GT(GS)_3_) was appended to the C-terminus of the wild-type protein and then connected to residue 1 of the wild-type sequence. Each circular permutant is identified by the wild-type residue that is the new N-terminus of the protein. For example, CP33 refers to the circular permutant starting at residue 33, and CP1 refers to the wild-type sequence with the added C-terminal GS linker.

To assay the effect of circular permutation on folding, function, and the propensity to aggregate, we developed a simple gel-based assay to monitor the function and solubility of all circular permutants of HaloTag directly in *E. coli* cell lysate (Supp. Fig. 1b). We took advantage of the fact that HaloTag is an engineered haloalkane dehalogenase that covalently binds halogenated ligands (Fig. 1a), and therefore ligand binding can be monitored as a readout of protein function.

Ligand binding was assayed via staining using the fluorescent HaloTag ligand TMR (Promega^TM^). *E. coli* cells expressing individual circular permutations were lysed and samples of whole-cell lysate and the clarified supernatant were stained with TMR prior to being run on an SDS-PAGE gel. Gels were then imaged for TMR fluorescence to determine if a circular permutant is functional in the soluble fraction. These same gels were then stained with Coomassie to compare the relative amounts of protein present in each fraction (Fig. 1b, Supp. Fig. 1c-f). While several CPs (e.g. CP217, Fig. 1b) showed high solubility and functional levels, some functional CPs were aggregation-prone in cells. For example, CP33 is less abundant in the soluble fraction than in the whole cell lysate, but protein in the soluble fraction can nevertheless bind TMR (Fig. 1b). Other CPs such as CP36 and CP39 are nonfunctional, and show different levels of aggregation-propensity (Fig. 1b).

Function and solubility scores were then assigned based on gel assays performed for every circular permutant (Fig. 1c-e). Circular permutants showed a range of solubility and functional levels, with 180 CPs retaining a high level of TMR binding and another 53 CPs showing low levels of TMR-binding (see Supp. Fig. 2a, CP44 vs CP45) – thus 78% of HaloTag CPs can function in *E. coli* cell lysate (Fig. 1c). Interestingly, when cells were lysed with a detergent-based protein extraction reagent (BugBuster), 118 CPs showed a decrease in function and change in category, suggesting the detergents present in this lysis buffer could interfere with proper TMR-binding (Supp. Fig. 2-3). 90 CPs are considered highly aggregation-prone (at least threefold less-soluble than CP1, Fig. 1d). Interestingly, 48 of these aggregation-prone CPs are functional in freeze-thaw lysis conditions, while 25 are functional in detergent-based lysis conditions. In sum, these results show that circular permutation can differentially affect both solubility and function of HaloTag in *E. coli*.

Wild-type HaloTag produces more soluble protein when folding co-translationally than when refolding *in vitro* (33). To test whether CPs could fold more efficiently during refolding than when expressed in cells, we selected aggregation-prone CPs to determine whether functional protein could be purified from inclusion bodies (Supp. Fig. 4a). We first chose CP119 and CP277, which were both capable of binding TMR in gel experiments, but show low levels of soluble expression in *E. coli* cell lysate. Cells expressing CPs 119 and 277 were lysed and, after a hard spin, the insoluble pellet was isolated (Supp. Fig. 4a). The pellet was resuspended in buffer containing 8M urea, then diluted to initiate refolding and incubated overnight at 37 °C. Samples were then stained with TMR and run via SDS-PAGE. While CP119 could not solubly refold in these conditions, CP277 did refold solubly from inclusion bodies and could then bind TMR (Supp. Fig. 4b-c). Therefore, our assay indicates that CP119 only folds efficiently during co-translational folding, while CP277 folds potentially more-efficiently during refolding than during co-translational folding. This experiment was repeated with 32 other aggregation-prone CPs: four proteins could refold from inclusion bodies and bind TMR (Supp. Fig. 4c-d, blue), four produced soluble protein that could not bind TMR (Supp. Fig. 4c-d, orange), and the other 24 did not refold solubly (Supp. Fig. 4c-d, black).

To assay levels of protein function in living cells using a high-throughput approach, which then enables an easy assay of protein function in multiple cell types, we screened the HaloTag CP library in *E. coli* BL21 (DE3) cells (the same strain as used in gel analyses) via FACS-seq. Cells were transformed with the CP library, stained with TMR, and sorted via FACS based on in-cell TMR fluorescence (Supp. Fig. 5a-b). High-fluorescence and low-fluorescence cell populations were collected, and DNA was isolated from sorted and unsorted cells immediately after FACS. Samples were then prepared for deep sequencing with an Illumina MiSeq and sequencing data for each replicate were compared to determine data quality (Supp. Fig. 5c-d).

We then sought to identify which CPs were functional in each cell type by calculating enrichment scores for the high-fluorescence cell populations. It is important to note that these FACS-seq readouts only report on the overall level of TMR fluorescence and cannot distinguish subtilties that arise from differences in soluble protein levels versus differences in protein function, unlike our gel-based assays. Therefore, these functional scores represent a combination of protein function, solubility, and expression levels in cells. We used the DESeq2 RNA-sequencing analysis package, which calculates log-fold changes between the unsorted and sorted populations and makes score adjustments based on the depth of the sequencing reads for each variant (37). This package also calculates standard error and statistical significance values, which enables us to determine which CPs are significantly enriched for function in each FACS-seq dataset. Fig. 2a shows the calculated enrichment scores for each CP, with yellow boxes drawn to mark contiguous stretches of functional CPs in the BL21 FACS-seq dataset; CPs with scores greater than zero are considered functional. As anticipated, we see agreement between our *in vitro* gel assay functional scores and the enrichment scores calculated from our BL21 FACS-seq dataset (Fig. 2b, Pearson 0.52 < r < 0.54). Interestingly, upon examining the CPs that lose function in the detergent-based lysis gel assay dataset, we find that many of these CPs are also nonfunctional in the BL21 FACS dataset (Supp. Fig. 3). This suggests that the presence of detergent-like molecules or lipids in the cell may disrupt TMR binding and contribute to these negative FACS-seq scores.

**Figure 2.**
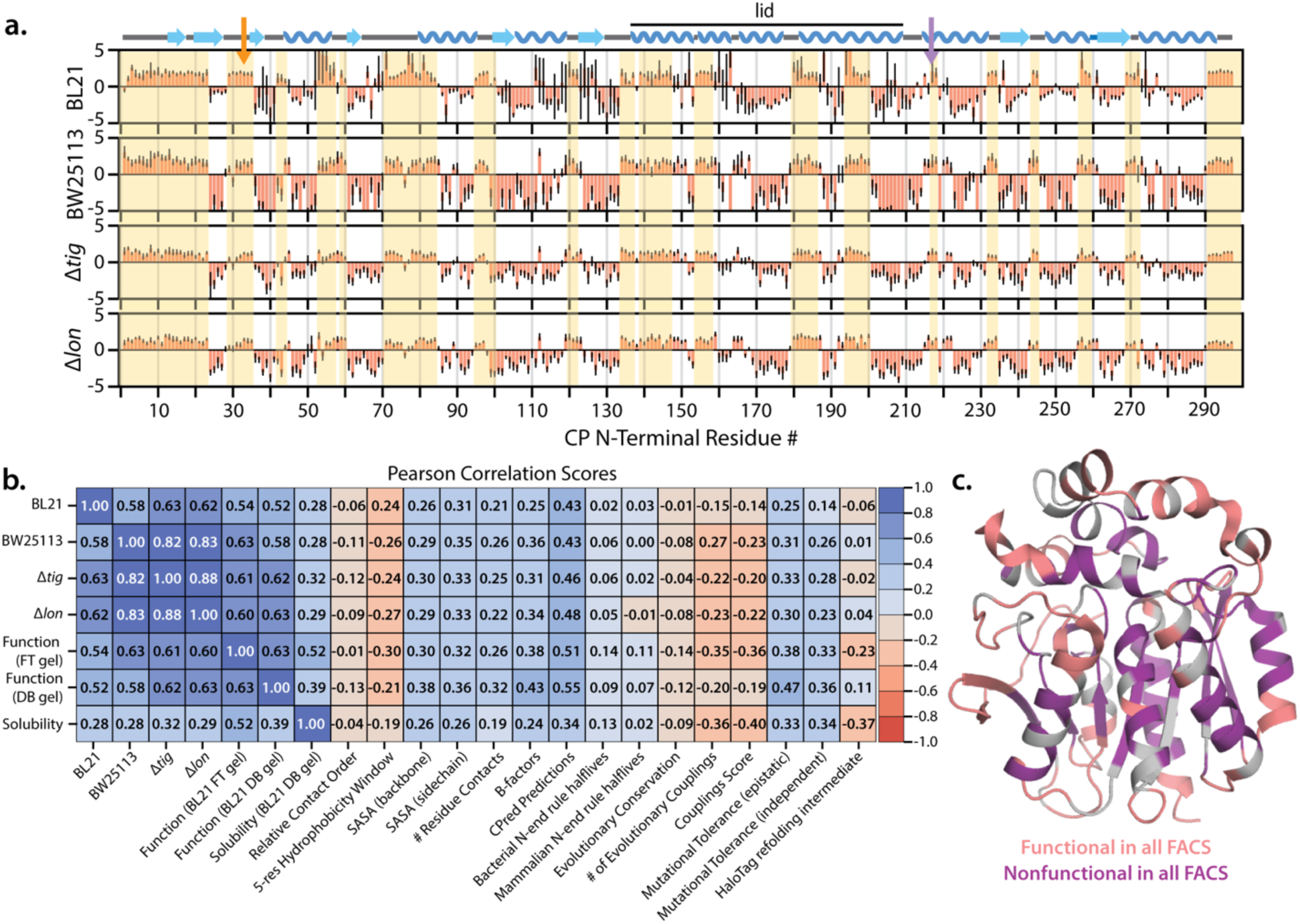
FACS-seq data for the CP library in different *E. coli* strains and correlations between functional scores and structural parameters. a. FACS-seq log-fold change enrichment scores for the CP library screened in BL21, BW25113, Δ*tig*, and Δ*lon* cells. Enrichment scores are plotted on a log_2_ scale and error bars show the score ± the standard error obtained from DESeq2 analyses. CPs with scores >0 are considered functional. Yellow boxes are drawn based on contiguous stretches of functional CPs in the BL21 dataset. Orange and purple arrows indicate positions of CP33 and CP217, respectively. b. Correlations between FACS, gel scores, and structural parameters. CPs with high levels of function are likely to have termini introduced in flexible, solvent-accessible, and hydrophilic regions of the protein. Functional CPs are also likely to have termini introduced in regions with high mutational tolerance and low degrees of evolutionary couplings based on an EVCouplings analysis. c. Summary of CPs robustly showing high- or low-function across all FACS datasets. CPs with consistent scores >0 are shown in pink, while CPs with scores <0 are shown in purple.

### CP function is highly robust in multiple cellular environments

The above high-throughput assay allows us to examine the effect of the cellular environment and specific proteostasis factors on the folding and function of each circular permutant. Because circular permutation changes which region of a protein is synthesized first during translation, we expected that CP nascent chains could differentially interact with the co-translational chaperone trigger factor, potentially impacting functional levels in cells. We were also curious to see whether abundance of Lon protease in the cell could impact the function of CPs prone to misfolding or aggregation, as Lon is deleted in BL21 expression strains and is therefore absent in the cellular environment where we carried out our gel assays. We therefore screened our library of HaloTag CPs in three additional cellular environments: the Keio collection trigger factor- and Lon-deletion strains (Δ*tig and* Δ*lon* respectively), in addition to the Keio parent *E. coli* strain BW25113 (38). Figure 2 shows that many CPs retain robustly high (Fig 2c, pink residues) or low (Fig. 2c, purple residues) levels of function across all cell types. When comparing these CP functional data in the Δ*tig* and Δ*lon* deletion strains with the BW25113 parent strain, we identify subsets of CPs that show changes in function in these altered cellular environments. However, these changes are not statistically significant (*p*>0.05) based on the DESeq2 tests comparing the input set of DNA sequences to the sorted pool (Supp. Fig. 6). Thus, perhaps surprisingly, this analysis suggests that the *in vivo* determinants of functional protein folding after relocating the N-terminus in HaloTag are independent of these cellular factors.

### High levels of CP function and solubility are correlated with termini insertion in flexible, solvent-accessible regions of HaloTag

Figure 2c shows that in general, highly-functional CPs have their termini introduced in lid loops, lid helices, core loops, and in scattered positions in core helices. Poorly-functional CPs have termini introduced in the core β-sheet and some core helices, though β-strand 1 and part of β-strand 2 tolerate termini insertion. The region of the lid α-helix H’ that packs together the core and the top of the lid is also intolerant to termini insertion, and it is likely that the introduction of flexible termini in this region impairs TMR binding.

With these qualitative observations, we set out to look for correlations between CP termini locations and specific structural parameters. We find that high levels of function (as determined by either our FACS or gel-assay scores) are correlated with termini insertion in regions with low hydrophobicity, high solvent-accessible surface area (SASA), and high crystal structure B-factors (Fig. 2b, Pearson r of 0.21 < r < 0.43). There are also similar correlations (0.19 < r < 0.26) between these parameters and high levels of soluble protein expression in *E. coli*. No correlation is observed between high function and relative contact order (RCO), which quantifies the average sequence separation between residues in contact in the native state. RCO has previously been shown to have strong correlations with changes in folding rate in small two-state refolding proteins, but not in circular permutants of a singular protein (39, 40). Based on our data with HaloTag CPs, RCO also does not predict high function or high solubility. We also find no correlations between high function and solubility of CPs and the N-end rule (−0.01 < r < 0.14), which describes how the N-terminal residue identity of a protein can influence the length of time a protein remains in the cell before degradation (41–43). Many of these structural parameters were also incorporated into the CPred machine learning model for predicting functional CP sites within a protein (44, 45). The CPred predictions for CP sites in HaloTag show good agreement with our identified functional CP locations (0.43 < r < 0.55).

We were also interested in studying how the observed changes in protein function for each circular permutant could relate to the evolutionary conservation of the new N-terminal residue. We used EVCouplings (v2.evcouplings.org, (46)) to calculate the sequence conservation and mutational tolerance of each residue in HaloTag based on an alignment with 4850 closely-related protein sequences. We theorized that termini insertion may be unfavorable in regions of the protein that are highly conserved or show low mutational tolerance. Protein regions under high evolutionary selection may be critical for proper structure formation and catalytic function, and thus would be incapable of accommodating insertion of flexible, charged termini through circular permutation. We therefore compared our FACS and gel assay scores to several parameters: sequence conservation at each position, the number of evolutionary couplings between residues (coupling scores reflect the likelihood that two residues are in contact and coevolving), and the mutational tolerance of each position calculated using two different models.

Residues with the highest evolutionary coupling scores are found primarily in the core β-sheet, as do the residues with the lowest mutational tolerance (Supp. Fig. 7). This suggests there are strong evolutionary restraints placed on the HaloTag core, and maintaining residue contacts with the β-sheet could be critical for folding. When compared with CP function based on our FACS and gel datasets, there is no correlation (−0.14 < r < −0.01) between high levels of CP function and high levels of sequence conservation, while there are stronger correlations (0.19 < r < 0.40) between low coupling scores/high mutational tolerance and CP function, as assessed by solubility and function (Fig. 2c). Overall, HaloTag tolerates termini insertion in flexible, surface-exposed regions of the protein where there are few contacts between residue side chains and low levels of evolutionary coupling to other residues in the protein.

### Biophysical analyses: circular permutation affects structure and stability

To probe the details of how permutation can affect the folding trajectory and energetics of a protein, we selected two circular permutants, CP33 and CP217, for detailed biophysical characterization *in vitro*. Both of these CPs have high functional scores in our FACS datasets and are topologically interesting. In our gel assay, CP217 is soluble and functional, while CP33 is functional but aggregation-prone. CP217 has termini introduced between the core and lid subdomains of HaloTag, and therefore contains the intact protein core N-terminal to the protein lid (Fig. 1b, light purple spheres). CP33 places the termini between β2 and β3, therefore moving β1 and β2 to the C-terminus (Fig. 1b, orange spheres). This could slow the formation and collapse of the core β-sheet during both refolding and co-translational folding.

To evaluate the overall structure of the permutants, we first turned to circular dichroism (CD) spectroscopy. Figure 3a shows the far-UV CD spectra of purified HaloTag, CP1, CP33, and CP217. With the exception of CP217, all show similar CD spectra. CP217, however, displays a slightly altered CD spectrum, suggesting CP217 contains less secondary structure than the other proteins.

**Figure 3.**
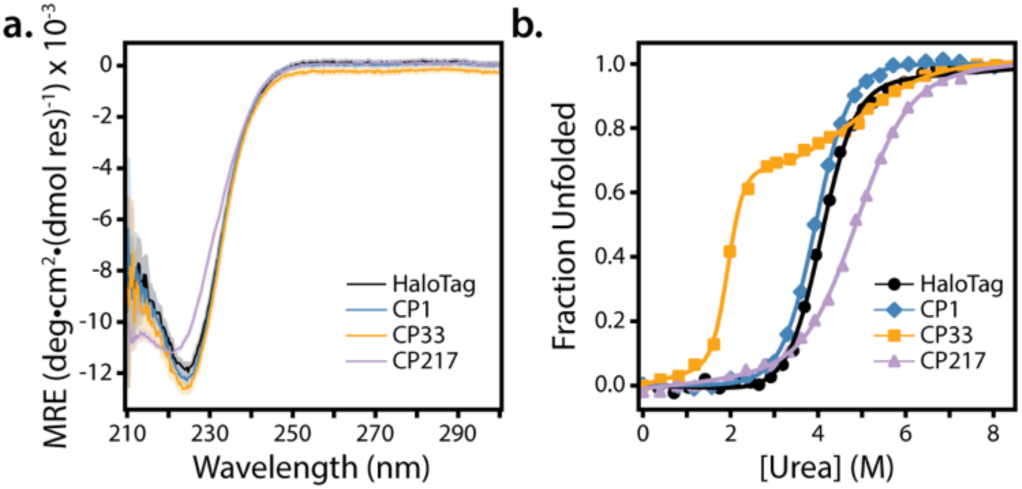
CD spectra and urea denaturation of CP variants. a. Far-UV CD spectra of HaloTag (black), CP1 (blue), CP33 (orange), and CP217 (purple). CP217 shows a decrease in CD signal relative to HaloTag. CD signal was averaged for 5 sec and measured every 0.25nm between 210 and 300nm. Shading represents the standard deviation of each measurement. b. Urea-induced denaturation monitored by CD at 225nm and fit with two- or three-state models. HaloTag (black circles) and CP1 (blue diamonds) show one unfolding transition around 4M urea, CP217 (purple triangles) shows one broad unfolding transition at 5M urea, and CP33 (orange squares) shows two transitions at 2M and 5M urea. CP33 populates an equilibrium intermediate not observed in HaloTag.

Protein stability was evaluated by equilibrium urea-induced denaturation, monitoring the CD signal at 225 nm (Figure 3b). These resulting denaturation curves were analyzed using a standard two-state (unfolded ⇌ folded) or three-state (unfolded ⇌ intermediate ⇌ folded) linear-extrapolation model (47, 48). With the exception of CP33, all show a single cooperative unfolding transition. The unfolding transitions for HaloTag and CP1 overlap with a resulting C_m_ of 4M urea (Table 1). CP217 shows a broader transition with a lower *m*-value. This lower m-value, along with the change in the CD spectrum, is consistent with a partially-unfolded native structure that has a smaller change in surface area upon unfolding. Interestingly, CP33 shows two transitions at 2M and 5M urea, uncovering an equilibrium intermediate not observed in the other proteins. After this initial analysis, we evaluated additional lid CPs, uncovering several CPs which show three-state unfolding with an early transition between 0-2M urea and a second transition observed at 5M urea (Supp. Fig. 8, Supp. Table 1). These particular CPs were selected based on the results of the GFP-domain insertion studies (CP156, 195) (49), a previously reported circular permutant (CP 143) (50), and on unusual results from our high-throughput studies (CP174 showed high function in gel assays but low function in FACS datasets). In sum, these data show the unfolding of many HaloTag CPs is less-cooperative than the unfolding of WT HaloTag, particularly those with termini insertion in the lid region.

**Table 1.** CD melt and kinetic refolding fit parameters. Errors represent the standard deviation of the fit.

|  | HaloTag | CP1 | CP33 | CP217 |
| --- | --- | --- | --- | --- |
| $\Delta G_{1,\text{fold}}$ (kcal/mol) | $-6.46 \pm 0.43$ | $-6.21 \pm 0.21$ | $-5.71 \pm 0.32$ | $-5.21 \pm 0.22$ |
| $m_1$ (kcal/mol•M) | $1.59 \pm 0.11$ | $1.58 \pm 0.05$ | $2.99 \pm 0.30$ | $1.06 \pm 0.05$ |
| $C_{m,1}$ (M) | $4.06 \pm 0.54$ | $3.93 \pm 0.26$ | $1.91 \pm 0.02$ | $4.92 \pm 0.27$ |
| $\Delta G_{2,\text{fold}}$ (kcal/mol) | - | - | $-3.78 \pm 0.60$ | - |
| $m_2$ (kcal/mol•M) | - | - | $0.76 \pm 0.21$ | - |
| $C_{m,2}$ (M) | - | - | $4.97 \pm 0.39$ | - |
| $t_{1/2,\text{fast,CD},10\text{C}}$ (s) | $611 \pm 9$ | - | $1362 \pm 22$ | $220 \pm 10$ |
| $t_{1/2,\text{slow,CD},10\text{C}}$ (s) | $4548 \pm 257$ | - | $12415 \pm 127$ | $1450 \pm 72$ |
| $A_{\text{burst,CD},10\text{C}}$ (MRE $\times 10^{-3}$) | $5.48 \pm 0.06$ | - | $4.50 \pm 0.02$ | $6.02 \pm 0.05$ |
| $A_{\text{fast,CD},10\text{C}}$ (MRE $\times 10^{-3}$) | $3.34 \pm 0.04$ | - | $1.12 \pm 0.01$ | $1.06 \pm 0.03$ |
| $A_{\text{slow,CD},10\text{C}}$ (MRE $\times 10^{-3}$) | $1.90 \pm 0.02$ | - | $1.92 \pm 0.01$ | $0.55 \pm 0.02$ |

### Amide hydrogen exchange monitored by mass spectrometry shows circular permutation can alter local energetics

To evaluate the local structure and stability of circular permutants, we turned to hydrogen-deuterium exchange monitored by mass spectrometry (HDX/MS) (Fig. 4a). HDX/MS follows the exchange of amide protons with solvent deuterons as a function of time at the resolution of individual peptides. Slowing of exchange, or protection, results from stabilization of backbone hydrogens in hydrogen bonds and/or lack of solvent accessibility (51, 52). Using HDX/MS, we were able to monitor peptides throughout the entire protein (99.7% coverage, Fig. 4b), allowing us to map and compare the local structure/energetics for all the permutants studied. In the absence of urea, WT HaloTag and CP33 show similar exchange profiles with slowed exchange in the expected regions of secondary structure (Fig. 4c-d, all peptides), indicating a wild-type structure with a well-folded protein core and lid in CP33. To probe the structure of the equilibrium intermediate observed for CP33, we monitored amide exchange under conditions where the intermediate is populated (3.5M urea). Most peptides in CP33 were fully exchanged within the first timepoint, with only the peptides spanning β5-7 and three core helices hB, hC, and hI showing measurable protection from exchange (Fig. 4c peptides 81-90 and 114-128, summary in Fig. 4e, more peptides in Supp. Fig. 9). These data suggest the CP33 equilibrium intermediate is comprised of a minimal folded core, while the lid, β1-4, and core helices hA, hJ, hK, and hL are unfolded.

**Figure 4.**
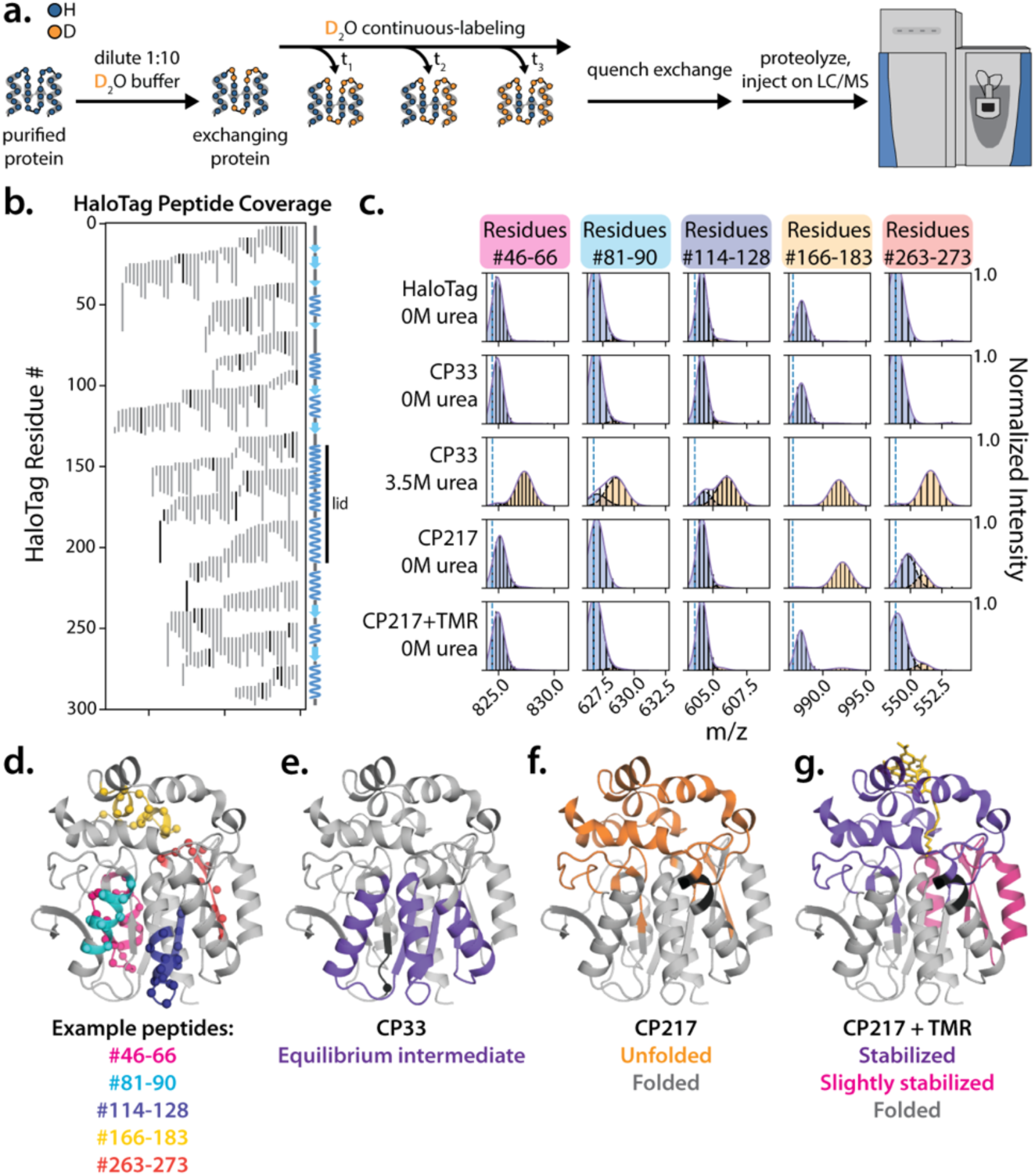
The CP33 equilibrium intermediate maps to the protein core, and the CP217 lid is stabilized upon binding of TMR. a. Schematic outlining continuous-labeling HDX/MS experiments. Protein is diluted into deuterated buffer, and samples are removed at various time points to quench hydrogen exchange. Samples are injected onto an LC/MS setup for inline proteolysis, peptide separation, and injection into the MS. b. Peptide coverage of HaloTag obtained from tandem MS/MS experiments. Peptides covering 99.7% of the HaloTag sequence are obtained. Black lines indicate the subset of peptides analyzed in continuous- and pulsed-labeling experiments. c. Mass spectra of five example peptides after 10s of HDX. Dashed blue line marks the monoisotopic mass of the peptide. Bimodal mass spectra are fit with a sum of two Gaussian curves (solid colored line), with the single-Gaussian fits represented in orange and blue (dashed black lines). Less-deuterated blue peaks correspond to folded protein, while the heavier orange peaks correspond to unfolded protein. Purified HaloTag and CP33 show similar protection from HDX in native conditions. When CP33 is equilibrated in 3.5M urea (where only the equilibrium intermediate should show structure), peptides either show high levels of exchange (are unfolded) or show a mixture of folded and unfolded populations (structured in intermediate). CP217 shows protection from HDX in the core, while the lid region is fully-exchanged and unfolded (166-183). CP217 lid peptides show protection from HDX when TMR is bound, in addition to stabilization of some core peptides (166-183, 263-273 respectively) d. Example peptides from c. mapped onto the HaloTag structure. e. Summary of all peptides structured in the CP33 intermediate (purple). f. Summary of peptides unfolded in CP217 (orange), and g. peptides that fold upon binding of TMR in CP217 (purple, highly-stabilized. Pink, slightly stabilized). Black residues in f-g do not have peptide coverage in CP33 or CP217, respectively.

In contrast to WT HaloTag and CP33, the native state of CP217 only shows protection from exchange in the core, with a highly-destabilized or unfolded lid that is fully-exchanged even in the absence of urea (Fig. 4c peptide 166-183, summary in Fig. 4f, more peptides in Supp. Fig. 9). These data suggest that the helical lid needs to be conformationally restrained at both its N- and C-termini in order to form a stable structure. Because our gel-based assays indicate that CP217 can fold and function (bind TMR), we carried out the same HDX/MS experiment on CP217 in the presence of the ligand TMR. Under these conditions, we see slowed hydrogen exchange in the lid and stabilization in some core peptides (Fig. 4c peptides 166-183 and 263-273, summary in Fig. 4g, more peptides in Supp. Fig. 9) consistent with folding upon ligand binding.

### Circular permutation impacts protein folding kinetics

The *in vitro* refolding of HaloTag is known to be aggregation-prone when carried out at 0.8M urea (37 °C), forming visible precipitate (33). CP33 and CP217 also show aggregation in these conditions, though with less apparent insoluble aggregate (Supp. Fig. 10). However, when refolded at 10 °C, all three proteins fold solubly and show burst, fast, and slow-folding kinetic phases as monitored by CD (Fig. 5, Supp. Fig. 10). HaloTag refolds with a t_1/2, fast_ of 6.1 x 10^2^ s and a t_1/2, slow_ of 4.5 x 10^3^ s (Table 1). CP33 refolds slightly slower, with fast and slow refolding t_1/2_s of 1.3 x 10^3^ s and 1.2 x 10^4^ s respectively, while CP217 refolds faster than HaloTag with fast and slow refolding t_1/2_s of 2.2 x 10^2^ s and 1.4 x 10^3^ s respectively (Table 1). The amplitude of the burst phase is similar in each CP suggesting all might fold through a similar burst phase intermediate (Table 1). In CP217, most of the CD signal is obtained during the burst phase, likely due to the lack of stabilization observed in the protein lid (see below).

**Figure 5.**
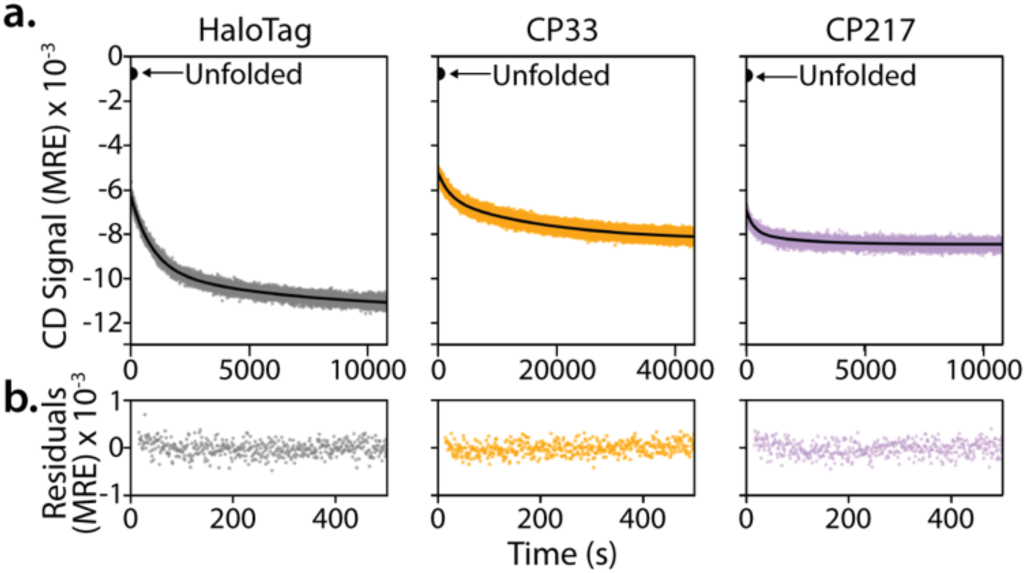
HaloTag CPs show altered refolding kinetics. a. Refolding to 0.8M urea monitored by CD at 225nm for HaloTag (gray), CP33 (orange), and CP217 (purple) at 10 °C. Refolding traces are fitted with biphasic kinetics (black lines), and CD mean residue ellipticity (MRE, (deg•cm^2^•(dmol•res)-1) x 10^-3^) of the unfolded protein is marked with a black point and black arrow. b. Fit residuals for the first 500s of refolding time.

### CP33 populates a smaller refolding intermediate than HaloTag, which resembles its equilibrium intermediate

The changes in aggregation-propensity between the different permutants suggest there may be a change in the folding pathway. To investigate the structural details of the folding trajectories, we turned to pulsed-labeling HDX/MS at 10 °C (33, 53, 54). Briefly, purified protein was denatured in high [urea], then rapidly diluted into low [urea] to initiate the refolding process. At various refolding timepoints, samples were further diluted into a deuterated buffer to initiate a ten-second pulse of hydrogen-deuterium exchange. Exchange was then quenched with ice-cold, low-pH, high-denaturant buffer, and samples were frozen until injected into our LC-MS setup (Fig. 6a). Bimodal mass spectra (heavy and light peaks) were observed for most peptides in all three permutants, where the heavier peak corresponds to the unfolded state and the lighter peak corresponds to a folded state protected from exchange (Fig. 6b-c). As the protein refolds, the heavier peak decreases in intensity and the lighter peak increases in intensity. These mass spectra were fitted to a sum of two Gaussian curves to calculate the population percentages in the folded/unfolded states at each timepoint (55).

**Figure 6.**
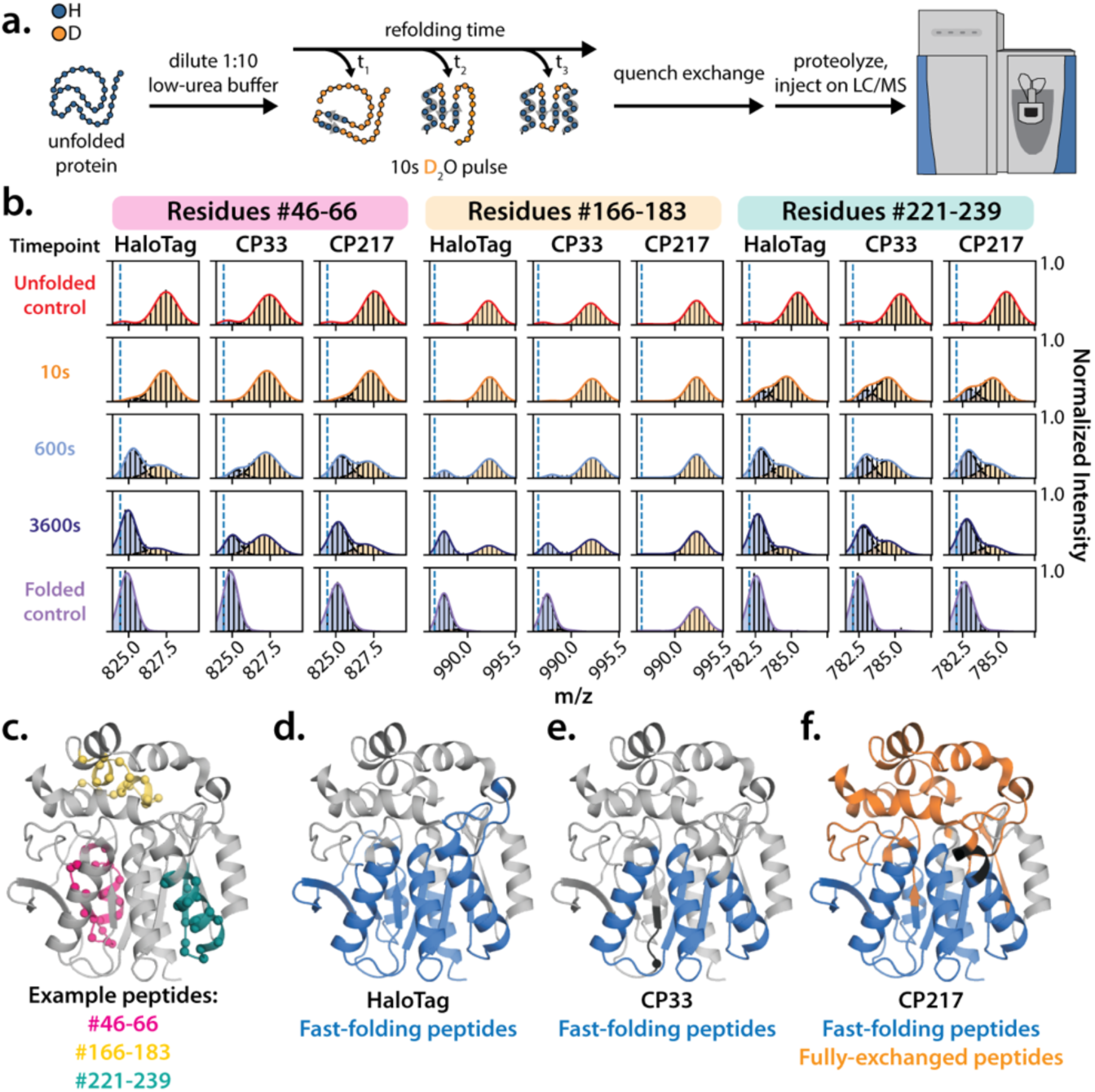
CP33 refolds through a different trajectory than HaloTag. a. Schematic outlining pulsed-labeling HDX/MS experiments. Protein denatured in 7.5M urea is diluted to 0.8M urea, and samples are removed at each refolding time point to dilute into deuterated buffer. Hydrogen exchange is quenched after 10s, and samples are injected onto an LC/MS setup for inline proteolysis, peptide separation, and injection into the MS. b. Mass spectra of three example peptides from HaloTag, CP33, and CP217 after 10s, 600s, and 3600s of refolding. Dashed blue line marks the monoisotopic mass of the peptide. Bimodal mass spectra are fit with a sum of two Gaussian curves (solid colored line), with the single-Gaussian fits represented in orange and blue (dashed black lines). Less-deuterated blue peaks correspond to folded protein, while the heavier orange peaks correspond to unfolded protein. Peptides are considered fast-folding if the measured populations are >50% folded at 600s. c. Example peptides from b. mapped onto the HaloTag structure. d-f. Summary of all fast-folding peptides in HaloTag, CP33, and CP217 are colored in blue. Gray regions are slow-folding, and f. orange regions in CP217 are fully-exchanged in the folded protein control. e-f. Black residues in CP33 and CP217 represent regions with no coverage due to termini insertion at a new location in each CP, and spheres indicate the new N-terminal residue in each CP.

We categorized peptides as fast-folding if they are >50% in the folded peak after 600s of refolding. This 600s time point was selected for analysis to correspond with the t_1/2_ of the HaloTag fast-folding phase obtained in CD experiments, and because several peptides show distinct differences in %folded at this time point (Fig. 6b-c). For example, in Fig. 6a, peptide 221-239 is >50% folded at 600s for all three proteins and is categorized as fast-folding. Peptide 166-183 is <50% folded at 600s, and is characterized as slow-folding in all three proteins. Finally, peptide 46-66 is >50% folded in HaloTag and in CP217, but >50% <u>un</u>folded in CP33. This peptide is therefore fast-folding in HaloTag and CP217 but slow-folding in CP33.

In WT HaloTag (Fig. 6d), we find that the fast-folding intermediate is made up of peptides spanning β1-7 in the core β-sheet and in five helices (hA, hB, hC, hI, and hL) packing onto this core sheet. In CP217, the same set of fast-folding peptides observed in WT HaloTag are also fast-folding in CP217 despite having a predominantly unfolded lid region in the native state (Fig. 6f). Remarkably, the early intermediate in CP33 is notably different than the others (Fig. 6e). The CP33 refolding intermediate is smaller than that observed in HaloTag, with a refolding core comprised of only β5-7 and the three core helices packing onto one side of this smaller sheet (hB, hC, and hI). β1-4 and helices hA and hL are now slow-folding. These fast-folding peptides are the same peptides that make up the CP33 equilibrium intermediate, indicating that CP33 folds through this stabilized equilibrium intermediate *in vitro*. This smaller fast-folding structure is also consistent with the smaller observed amplitude of the fast-folding phase in CD experiments (Table 1).

In addition to the fast-folding core, there are also a set of peptides that show a burst phase or gain of structure within the first ten seconds of refolding, defined as a leftward shift in the heavy unfolded peak (indicating protection) when comparing the unfolded control to the first ten-second time point (Supp. Fig. 11a-b). In Fig. 6a, peptide 221-239 shows a leftward shift in the unfolded population of 2.5 Da and peptide 46-66 shows a 0.5 Da leftward shift. By contrast, peptide 166-183 shows no shift and in fact no lid peptides show this burst activity.

In all three proteins studied, the burst peptides map to the core β-sheet and helices, suggesting an initial collapse of the core within the deadtime of the experiment. All HaloTag fast-folding peptides show this burst-phase behavior except for peptides 9-18 and 286-294 (Supp. Fig. 11c, the N-terminal β-strand and C-terminal helix). CP217 and CP33 show highly similar behavior to HaloTag with notable exceptions, suggesting changes in the folding landscape: in CP33, the same set of peptides that have burst-phase behavior in HaloTag also have burst-phase behavior in CP33 (except for peptide 19-32 which is located at the CP33 C-terminus) (Supp. Fig. 11a-b, summary in Supp. Fig. 11d). However, some of these peptides now fold slowly after the burst in CP33 (e.g. Fig. 6b peptide 46-66). Therefore, CP33 shows a change in folding pathway for these regions of the protein after the initial collapse occurs. In CP217, peptide 286-294 uniquely shows a burst-phase gain of 0.5 Da that is not observed in HaloTag or in any other CP analyzed with this approach (Supp. Fig. 11a-b, summary in Supp. Fig. 11e).

As an orthogonal approach to monitor the change in structure of the refolding intermediate in CP33, we turned to pulsed thiol-labeling experiments (33). This experiment monitors folding at the level of side-chain (cysteine) burial. HaloTag contains two native cysteines at positions 61 and 262. Based on the refolding intermediates identified using HDX/MS, C61 should gain structure and protection from solvent during the fast-folding phase in WT HaloTag while C262 should be slow-folding. Our model predicts that both cysteines should gain protection slowly in CP33. We monitored the extent of cysteine modification over time by pulse-labeling the refolding protein with a fluorescein-conjugated maleimide dye at different refolding timepoints, then quantifying the extent of labeling at each time point on an SDS-PAGE gel based on in-gel fluorescence (33). Rates of protection from thiol-labeling can therefore be measured to calculate site-specific rates of refolding (Supp. Fig. 12a-c). WT HaloTag shows biphasic protection from cysteine-labeling, with t_1/2_s for the fast- and slow-phases on the same orders of magnitude observed in CD experiments, consistent with only one cysteine (C61) gaining protection in the fast-folding phase. CP33, on the other hand, shows slow protection and single-phase kinetics, consistent with both cysteines being located in the slow-folding region of the protein. This slow rate of protection is also consistent with the slow-phase folding observed by CD (Supp. Fig. 12d-f, Supp. Table 2). Our findings that CP33 contains a change in folding trajectory are therefore supported by experiments probing both secondary (via HDX) and tertiary (via thiol labeling) structure.

## Discussion/Conclusions

We have taken a comprehensive approach to investigate how repositioning the N-terminus impacts a protein’s energy landscape by evaluating every possible circular permutant of the protein HaloTag. Relocation of protein termini to a new position through circular permutation has the potential to affect protein structure and function both *in vivo* and *in vitro*. For instance, placement of the N-terminus is expected to bias folding pathways and could either favor or disfavor aggregation-prone intermediates, which could lead to novel aggregation or engagement of proteostasis factors not essential for the wild-type protein. To evaluate these features, we examined the folding and function of all 297 circular permutants using both an *in vitro* gel assay and an in-cell high-throughput FACS screen. With our gel-based assay, we find that HaloTag is highly soluble when overexpressed in bacterial cells, yet some functional CPs show a novel propensity to aggregate, which is not observed in the wild-type protein. Termini insertion within the protein core is most likely to yield aggregation-prone proteins, while the lid is amenable to termini insertion while retaining both high solubility and functional levels. In general, HaloTag is very tolerant of circular permutation, with 60% of CPs retaining high levels of function and another 18% retaining low levels of TMR binding.

A combination of FACS and next-generation sequencing allowed us to screen every possible HaloTag CP in four strains of *E. coli*: the BL21 expression strain, the Keio collection parent strain BW25113, and the Keio Δ*tig* and Δ*lon* strains knocking out the co-translational chaperone trigger factor (TF) and the intracellular protease Lon (38). Since circular permutation inherently changes the *in vivo* order of synthesis of secondary structural elements, some CPs generate highly-hydrophobic stretches of residues at their N-termini as the protein core is synthesized. These hydrophobic regions might be selectively recognized by, and interact with, TF during synthesis, especially if the collapse of the hydrophobic core is delayed co-translationally. Additionally, TF contains a peptidyl-prolyl *cis/trans* isomerase domain (56), and could be required for the *in vivo* folding of HaloTag as it contains three *cis*-proline peptide bonds in the wild-type structure and proline makes up 10% of its sequence composition. Therefore, we expected to identify changes in functional protein folding in the Δ*tig* cellular environment. We also investigated the effect of Lon protease on CP function, as Lon is a highly-conserved intracellular protease responsible for clearing misfolded proteins that is deleted in BL21 cells (57). We were interested to see if any of the functional, aggregation-prone proteins identified in our gel-based assays would show changes in function in the presence or absence of this protease in BW25113 cells.

Contrary to our expectations, we found that the effect of circular permutation on HaloTag is robust to changes in the cellular proteostasis machinery. There are no significant changes in function for CPs when comparing the Δ*lon* and parent BW25113 datasets, and only one CP (CP290) becomes significantly nonfunctional in the absence of TF. After comparing various structural parameters to the gel-assay and FACS scores, we find that termini insertion in buried, restricted locations is most disruptive to HaloTag function and solubility when expressed in cells independent of the strain studied. While there are good correlations between these parameters and our CP scores, there still does not seem to be a perfect predictor of high levels of CP function.

What structural features are successful for starting a protein sequence? The CPred server applies several machine learning techniques to calculate the probability of successful termini insertion at each position in a protein (44, 45). Indeed, we see a strong correlation between our data and predictions of viable CP locations from CPred (Pearson r 0.43 < r < 0.55), though the correlation between CP solubility and CPred predictions is slightly worse (Pearson r = 0.34). Termini relocation to regions of the protein with low evolutionary coupling scores and high mutational tolerance is also more likely to yield a functional CP. It is interesting that termini insertion in these highly-coupled regions of the protein is deleterious to protein function and solubility. This could be due to the disruption of conserved stretches of contacts with the introduction of flexible, charged termini. These strong contacts could also be critical to proper formation of folding intermediates, and breaking these contacts through termini insertion could disrupt intermediates populated co-translationally.

Our results, therefore, suggest that a) the successful folding of HaloTag is highly-influenced by termini location, and that b) the misfolding and/or aggregation of CPs is impacted more by changes in folding and introduction of termini in conformationally-restrained regions than by changes in interactions with proteostasis factors. Functional CPs are most likely to have termini introduced in conformationally-flexible, surface-exposed regions with few to no side-chain contacts with other residues. Additionally, one of the strongest predictors of aggregation-propensity in CPs is termini insertion within the fast-refolding intermediate we identified by HDX/MS (Pearson r = 0.37). Termini insertion in this region likely disrupts the collapse and stabilization of the refolding core, which would increase the exposure of hydrophobic residues and the likelihood of forming intermolecular contacts leading to aggregate formation.

Interestingly, relocating the termini in HaloTag differentially impacts the energy landscape of the native state. The CPs we examined biophysically show differences in stability between the lid and core subdomains compared to the wild-type protein, where the folding of these regions is highly coupled. In fact, several CPs show three-state equilibrium behavior, in contrast to the two-state nature of WT HaloTag, and the lid region of the protein is also highly-destabilized in many CPs despite retaining high levels of function. This was initially surprising given that the lid makes up most of the ligand-interacting surface during binding, but given the aliphatic nature of HaloTag ligands, the lid likely collapses onto the long carbon chain in Halo-TMR.

In addition to the structure and stability of the folded state, the folding process itself is affected by circular permutation. Not surprisingly, the kinetics of folding depend on the specific permutation. What is perhaps more surprising, however, is that the folding pathway is altered, and here we identify a change in the structure of a refolding intermediate with circular permutation. The fast-refolding intermediate of CP33 comprises less of the core region than wild-type HaloTag. CP33 and CP217 also show less aggregation-propensity than HaloTag, though the mechanism of avoiding aggregation is unclear. Interestingly, CP33 is aggregation-prone when expressed in cells, suggesting there could also be a change in the co-translational folding pathway for this CP as compared to WT HaloTag. These changes in protein energetics, folding rates, and folding pathways are consistent with previous observations in circular permutant studies. For example, in a randomized CP study of DsbA, an equilibrium intermediate is observed in chemical denaturation experiments with an oxidized CP that is not observed in wild-type DsbA, similar to what we see in CP33 and some of the lid CPs of HaloTag (58). In CP studies of the ribosomal protein S6 from *T. thermophilus*, CPs show changes in protein stability and folding kinetics compared to wild-type protein (59). Finally, ɸ-value analyses on CPs of the small ribosomal protein S6 also present evidence for folding through parallel pathways. α-spectrin SH3 CPs show changes in their folding transition states, indicating that refolding through different trajectories may be a general feature of circular permutation (9, 59).

This work adds to the small database of comprehensive studies on the effects of circular permutation within a protein. In a systematic study of *E. coli* dihydrofolate reductase (DHFR) circular permutants (60), 10 contiguous stretches of CPs were identified that are incapable of folding, suggesting that these regions are critical “folding elements” in DHFR. These regions include 10 of the 13 residues known to gain early (6ms) protection during folding as determined by pulsed-labeling HDX-NMR (61). Similarly, in a randomized circular permutation-study examining 65 CP sites out of 189 possible in DsbA (58), functional CPs did not have termini introduced in certain helices and β-strands, and rationally-designed CPs in these regions were inactive. Our HaloTag FACS-seq data also identify contiguous stretches of nonfunctional CPs, many of which are found within the fast-folding HaloTag intermediate, showcasing how termini insertion within fast-folding structural elements is highly disruptive to proper folding and function.

Our systematic demonstration that termini relocation in HaloTag can drastically impact soluble, functional protein expression in a manner independent of changes in cellular environment has important implications for both *de novo* protein design and for engineering proteins for improved expression. They may also shed light on the changes in the folding of proteins proteolytically processed in the cell, resulting in a new N-terminus. We cannot speculate on how the trends we observe between successful termini relocation and structural parameters would extend to larger, more-complex protein systems. It would be interesting to perform similar CP studies and test our observations in proteins that can undergo changes in oligomeric state, regulation by a binding partner, or that are known chaperone substrates. These data could then be used to train additional CP prediction models or to help improve existing models, which to date have primarily been developed using a small number of systematic and randomized CP datasets (44, 45). Additionally, performing FACS-seq in cellular environments lacking other co-translational chaperones such as the DnaK/DnaJ/DnaE system or post-translational chaperones such as the Hsp90s, or growing cells at lower temperatures to decrease rates of translation in the cell, could provide further insight into whether these changes in environment are sufficient to modulate the successful folding of HaloTag CPs. We also recognize that all of our in-cell functional results come from experiments in which we are inducing overexpression of CPs with an IPTG-inducible promoter, and this could mask any subtle changes in protein function. Given that multiple chaperones often interact with the same protein clients, it is also possible that no effect is seen on CP function in the Δ*tig* results presented here due to compensating interactions with DnaK and other chaperones. It would therefore also be interesting to perform these experiments under a more tightly-regulated titratable promoter, or to examine changes in function between overexpression and leaky expression conditions, to see if reducing the amount of CP being made in each cell then impacts functional protein folding.

## Materials and Methods

### Building the tandem HaloTag vector and sub-cloning 298 Circular Permutants

A Tandem HaloTag plasmid containing the HaloTag-GTGSGSGS-HaloTag cDNA sequence was built via Gibson Assembly (NEB, (62)) of two HaloTag genes, the linker sequence, and a parent vector containing a pUC19 origin, KanR, and the T7 promoter/terminator sequences flanking the coding region (pNRD_BL21). Primers were then designed to amplify gene fragments from the Tandem HaloTag sequence for every possible CP, where CP1 (wild-type HaloTag) contains the coding sequence for HaloTag with a C-terminal linker sequence, and CP298 contains the intact linker sequence N-terminal to a HaloTag gene. Each primer pair also contains overhang sequences to introduce an AUG start codon and UAG stop codon flanking the CP gene, in addition to binding sites for the Type IIS restriction enzyme Esp3I (an isoschizomer of BsmBI active at 37 °C). 298 PCRs were done to sub-clone the gene for each individual CP from the Tandem HaloTag plasmid, and a small volume of each PCR was pooled (CPLib) based on the relative intensity of each PCR product assessed on a 1% agarose gel. The CPLib gene pool was then inserted into a pNRD_BL21 PCR fragment containing compatible Esp3I restriction enzyme sites using a homemade Golden Gate Assembly reaction (pCPLib_BL21, (63)).

### Building a glycerol stock library for every HaloTag CP

Commercially available *E. coli* BL21 cells (Sigma-Aldrich) were transformed with the pCPLib_BL21 vector library via electroporation. Small volumes of the outgrowths from the transformation recoveries were plated and grown out overnight, and 1.5mL cultures in 96-well blocks were inoculated with colonies from the plate the following evening. After 16hrs of growth shaking at 37 °C, 0.5mL of saturated cultures were removed and diluted 1:1 in 50% glycerol in another 96-well block to make glycerol stocks of each colony. Glycerol stocks were stored at −80 °C, while the remaining saturated cultures were miniprepped and Sanger sequenced at the UC Berkeley Sequencing Facility. Sequencing results were then used to identify which CP was present in each glycerol stock. This was done twice with the full library of CP sequences, twice with a sub-library containing CPs that had not yet been isolated, and the final 68 CP genes were individually cloned into the pNRD_BL21 backbone, transformed, and sequenced to make glycerol stocks of each individual CP.

### A gel-based assay for determining CP function and solubility

5mL overnight cultures were inoculated from the BL21 E. coli glycerol stocks for each CP. The following morning, fresh 5mL cultures were seeded from the saturated overnight, then allowed to grow for 3hr before inducing protein expression with 1mM IPTG. After 2hr, cells were pelleted and lysed. Lysis was performed for each protein using two different methods to avoid artifacts that could arise from the chosen lysis method. Lysis was done either by resuspending the cell pellet in 0.8mL of the detergent-based BugBuster Protein Extraction Reagent (Millipore) and agitating for 20min on a rotator, or by resuspending the cell pellet in 0.8mL of 2μg/mL lysozyme in 10mM Tris pH 7.5, 0.1M NaCl, and 1mM EDTA, then doing three cycles of snap freezing the lysate with liquid nitrogen then thawing the cells in a 37 °C water bath. 10μL samples of whole cell lysate (L) were taken, then the remaining lysate was spun at 11,000rpm for 10min to pellet the insoluble fraction. 10μL samples of the clarified supernatant (S) were obtained.

L and S samples were stained with 20μL of 2.5μM TMR ligand diluted in 25mM HEPES pH 7.5, 15mM Mg(OAc)_2_, 150mM KCl, and 0.1mM TCEP (HKMT buffer), for a final staining concentration of 1.67μM TMR. After staining for 20min, L and S samples were boiled for 5min at 95C in an SDS loading buffer. Boiled samples were mixed well, then 15uL of sample were loaded into a NuPAGE 4-12% Bis-Tris 1.5mm gel and run in an MES running buffer. Gels were imaged for TMR fluorescence using the TAMRA filter on a Typhoon gel scanner, then Coomassie stained for an hour before moved into a destaining solution. Destained gels were then imaged for Coomassie staining.

Functional scores for each CP were measured by dividing the fluorescence of the CP protein band in the S fraction by the fluorescence of a band in the ladder, then dividing that quantity by the intensity of the same S fraction CP band in the Coomassie image normalized by a band in the ladder for samples where freeze-thaw lysis was used). Functional scores were also determined for the detergent-based BugBuster lysis method.

Solubility scores for each CP were calculated based only on the Coomassie staining images of the gels. The intensity of the CP band in the S fraction sample was normalized to the intensity of the band directly above the CP, and this value was then divided by the intensity of the band in the L fraction normalized by the intensity of the band above the CP for samples where the detergent-based BugBuster lysis was used. Solubility scores were then normalized to the solubility of wild-type HaloTag (CP1).

### Building the pNRD_ASKA expression vector for screening the CP Library in *E. coli* BW25113 cells and in Keio collection strains

Given that the *E. coli* strains used in the Keio single-gene deletion library (38) do not contain T7 protein expression systems, a new expression vector was needed to allow for IPTG-based induction of expression of our CP Library. The expression vector used in the ASKA library paper (64) was thus obtained from the Arkin Lab at UC Berkeley. To prepare this vector for use in our FACS-seq experiments, a four-piece Gibson Assembly (62) was first performed to simultaneously remove four Esp3I restriction sites in the backbone (pNRD_ASKA). The backbone was then PCR amplified to add Esp3I restriction sites and sequences complementary to Golden Gate assembly (63) with the CPLib gene pool.

### Preparing the CP library for FACS-seq in BL21, BW25113, Δ*tig*, and Δ*lon* cells

The CPLib gene pool was cut and pasted into either the pNRD_BL21 expression vector or the pNRD_ASKA expression vector via Golden Gate Assembly (63) with T4 DNA ligase and the restriction enzyme Esp3I (NEB). Golden Gate reactions were then cleaned up using a Zymo Clean and Concentrator kit before electroporation.

All Keio collection strains were obtained from the Deutschbauer Lab at UC Berkeley. Homemade electrocompetent BW25113 and Keio collection strains were prepared before transformation with the CPLib_ASKA vector pool. Fresh 50mL cultures were seeded from saturated overnight cultures and grown to an OD_600_ of ~1.0 before pelleting the cells. Cells were washed three times with 50mL ice-cold sterile water, then resuspended to a final volume of ~100μL cold water. 2μL DNA were mixed with 25μL cells, then diluted with cold sterile water to 100μL before electroporation in an ice-cold 1mm cuvette (BioRad MicroPulser). Cells were then recovered in 900μL LB for 1hr shaking at 37 °C before removing 5uL for a plated dilution series to determine transformation efficiency. The remaining cells were then pelleted and resuspended in LB containing the appropriate selection antibiotic(s), and grown to saturation overnight. Commercial electrocompetent BL21 (DE3) cells (Sigma-Aldrich) were transformed with the CPLib_BL21 library following standard protocols.

Fresh 5mL cultures were inoculated the following morning from the saturated overnights if the transformation efficiency was greater than 10^6^ cfu (>3,000x the library size). Cells were grown for three hours before inducing protein expression with 1mM IPTG, then allowed to continue shaking at 37 °C for an additional hour. 200μL of cells were removed and stained with 5μM TMR for 1hr at room temperature. Cells were then washed three times with 1mL PBS buffer (137 mM NaCl, 2.7 mM KCl, 10 mM Na_2_HPO_4_, and 1.8 mM KH_2_PO_4_ pH 7.4) and diluted to 2mL total volume of cells before sorting.

### Screening the CP library in bacterial cells for function via FACS-seq

Stained cells were sorted on a Sony SH800S cell sorter calibrated with a 70μm chip when working with bacterial cells. For bacterial sorting experiments, gating was set based on a negative control cell population (e.g. cells containing a vector expressing RNaseH) and a cell population expressing wild-type HaloTag. At least 1E6 cells were collected at room temperature from the high-fluorescence and low-fluorescence gates for three replicates. All remaining unsorted cells in addition to the sorted cell populations were then miniprepped (Takara NucleoSpin Plasmid miniprep kit), and DNA was stored at 4 °C short-term or at −20 °C for long-term storage.

The sorted and unsorted DNA pools were then prepared for Illumina MiSeq next-generation sequencing. Illumina adapters were added onto each pool through two consecutive PCRs with 15 and 20 cycles to reduce PCR-introduced errors. PCRs were pooled based on relative intensity of each PCR product on a 1% agarose gel, then the pooled amplicon libraries were gel extracted and quantified using the QuantIt dsDNA assay kit (Invitrogen ref Q33130) or via qPCR at the Vincent J. Coates Genomics Sequencing Lab at UC Berkeley. Amplicon libraries were then denatured and loaded at 10pM with 10% PhiX on an Illumina MiSeq Nano chip to quantify the sample loading density in a 50-cycle run, and loading concentrations were then optimized for a 300-cycle sequencing run with a full MiSeq chip. Forward reads were then analyzed using an in-house Python script to count how many instances of each CP occurred in each sequencing pool. CP counts and enrichment scores were then calculated using in-house Python scripts. The DESeq RNA-sequencing analysis package was also used to determine enrichment scores for each CP, to measure error between replicates, and to assign p-values to each score (37).

### Computational analyses of HaloTag structural parameters

Relative contact order (RCO) calculations were done with the TMR-bound HaloTag structure (PDB 6u32). A Python script was written to make a pseudo-PDB file for each CP sequence, and the RCO was calculated for each PDB file using a Python script adapted from the pdbtools GitHub repository (https://github.com/harmslab/pdbtools). Residues with an atom within 6Å of another residue are considered in close contact for these calculations. Backbone and side chain solvent-accessible surface area (SASA) were calculated for each residue with a Python script using a Shrake-Rupley algorithm from the BioPython package (65, 66). Hydrophobicity of residues surrounding the CP termini were calculated by Willow Coyote-Maestas (67). EVCouplings analyses were performed using the v2.evcouplings.org server and their default settings (46). CPred predictions were made using the CPred server at http://sarst.life.nthu.edu.tw/Cpred (45). B-factors are from the apo-HaloTag structure (PDB: 5uy1). N-end rule half-lives are as published (41–43).

### Protein expression and purification

Wild-type HaloTag and CPs were expressed in BL21 (DE3) cells in 1L cultures. Cells were grown to an OD600 ~0.6 then expression was induced with 1mM IPTG. Cells shook at 37 °C for three hours before pelleting cells and resuspending in 20mL of 50mM Tris pH 7.5. Pellet resuspensions were stored at −80 °C.

Thawed pellets were sonicated on ice to lyse cells, and clarified supernatant was filtered and loaded onto a 5mL HiTrap Q HP column (Cytiva 17115401). Protein was eluted off the column with a high salt gradient from 50mM Tris pH 7.5 to 50mM Tris pH 7.5, 1M NaCl. Fractions containing HaloTag were pooled and run over a HiLoad 16/600 Superdex 75 size-exclusion chromatography column (GE), buffer-exchanging into 25mM HEPES pH 7.5, 15mM Mg(OAc)_2_, 150mM KCl, and 0.1mM TCEP (HKMT buffer). Protein was immediately used for experiments, or stored short-term at 4 °C for use in experiments.

### Circular dichroism spectroscopy

CD experiments were done on either an Aviv 410 or an Aviv 430 CD spectrometer. For taking CD spectra, protein was prepared at 0.5mg/mL and equilibrated overnight at 37 °C in HKMT buffer. Far-UV spectra were measured by taking the average of five seconds of CD signal every 0.25nm from 210nm to 300nm in a 1mm quartz cuvette (Starna) at 37 °C. Spectra were buffer-corrected by subtracting the CD signal of the buffer at each wavelength, then converted to mean residue ellipticity using the following equation (68):

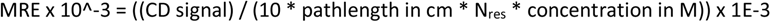

### Equilibrium denaturation experiments by CD

Chemical denaturation melts were set up by making two stocks of 0.1 mg/mL protein in HKMT and in HKMT with 8M urea, then mixing the 0M urea and 8M urea protein stocks at different ratios to make a range of final urea concentrations. Samples were equilibrated overnight at 37 °C, and the CD signal was averaged over 60 s. for each urea concentration at 225nm in a 0.5cm quartz cuvette (Starna) while stirring with a magnetic stir bar. Samples were recovered from the cuvette, and the actual urea concentration of each sample was determined using a refractometer. Melts were fit with Python scripts with the following global fit equations for two-state and three-state transitions, where F is the folded baseline, I is the intermediate baseline, and U is the unfolded baseline:

Two state melt (47):

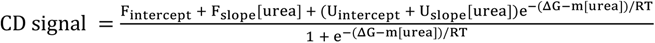

Three-state melt (48):

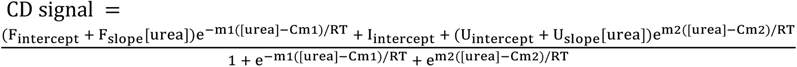

### Protein refolding by CD

Unfolded protein stocks were prepared at 2mg/mL in HKMT with 7.5M urea and equilibrated overnight at either 37 °C or 10 °C. HKMT refolding buffers were also prepared and equilibrated overnight with the appropriate amount of urea so that the final urea concentrations of refolding protein would be 0.8M or 7.5M urea for the unfolded control. For each experiment, 1.35mL of refolding buffer was placed in a 0.5cm cuvette in the CD, and the CD signal of the buffer was measured by averaging the CD signal for 60 s. at 225nm and the appropriate temperature. To initiate refolding, 0.15mL of protein was added to the same cuvette with refolding buffer, rapidly mixed up and down with a transfer pipette, capped, and placed in the CD while stirring with a magnetic stir bar. CD signal was measured every second for between 1hr and 12hrs (depending on the protein). The deadtime of mixing and starting the experiment was also recorded. Protein samples were recovered, and the final urea concentration was measured with a refractometer.

UV-Vis absorbance spectra were taken of the recovered samples from 240nm to 500nm and baseline corrected with spectra of the appropriate refolding buffer (Agilent Cary UV-Vis Compact Peltier). If light absorbance was observed between 300nm and 400nm, indicative of protein aggregate formation, the sample was spun at 16,000rpm for 10min. Spectra of the supernatants were then taken again to determine what concentration of protein remained soluble after refolding.

CD kinetic refolding traces were buffer-corrected and, if no aggregation was observed, traces were fit with single-exponential or double-exponential kinetics as done previously to measure rates of refolding (33).

### Continuous-labeling hydrogen-deuterium exchange

All HDX experiments were done with deuterated HKMT buffers containing zero or various concentrations of urea. Buffers were lyophilized and resuspended in D_2_O (Sigma-Aldrich 151882). Deuterated buffers were then aliquoted and stored in cryotubes at −80 °C.

To prepare proteins for HDX, 10uM HaloTag (HKMT, 0M urea) and 10uM CP33 (HKMT, 0M or 3.5M urea) were equilibrated overnight at 37 °C to mimic conditions used in CD melt experiments. For CP217 experiments with and without TMR, 10uM CP217 was equilibrated with 50uM TMR at 37 °C for 30min before equilibrating overnight at 10 °C to mimic pulsed-labeling HDX/MS conditions where the lid was first identified as being unfolded. To initiate exchange, samples were diluted tenfold into deuterated buffer. At each timepoint, the exchange mixture was diluted 1:1 into an ice cold 2x quench buffer (3.5M GdmCl, 1.5M glycine, 0.5M TCEP pH 2.4), flash-frozen in liquid nitrogen, and stored at −80 °C until LC-MS injection.

### Pulsed-labeling hydrogen-deuterium exchange

For the 7.5M urea buffers used in pulsed-labeling experiments, buffers were lyophilized and deuterated three times to ensure full deuteration of the urea. Deuterated buffers were then aliquoted and stored in cryotubes at −80 °C.

To prepare samples for refolding, 50uM protein in HKMT with 7.5M urea was equilibrated overnight at 10 °C and then diluted tenfold into a temperature-equilibrated refolding buffer to make the final refolding urea concentration 0.8M. At each folding time point, the refolding protein solution was diluted tenfold again into deuterated HKMT with 0.8M urea and then quenched after 10s of exchange by dilution 1:1 into an ice cold 2x quench buffer (3.5M GdmCl, 1.5M glycine, 0.5M TCEP pH 2.4, flash frozen in liquid nitrogen, and stored at −80 °C until LC-MS injection. Unfolded controls were prepared by diluting unfolded protein equilibrated with 7.5M urea into a 7.5M urea mock “refolding” buffer, and were pulsed with deuterated 7.5M urea HKMT following the same procedure as the refolding samples. Folded controls were prepared by diluting protein equilibrated at 0.8M urea into a 0.8M urea refolding buffer, then pulse-labeled with deuterated 0.8M urea HKMT buffer and then following the same procedure as the refolding samples.

### Protease digestion, LC-MS, and peptide identification

Digestion and LC-MS was performed as previously described (55). Briefly, samples were thawed immediately before injection into our LC-MS setup, which is comprised of a Trajan LEAP valve box connected to an Ultimate 3000 LC and Q-Exactive Orbitrap MS. Inline digestion was performed with porcine pepsin (Sigma-Aldrich P6887) conjugated to beads (Thermo Scientific POROS 20 Al aldehyde activated resin 1602906) and packed into a homemade protease column, and samples were desalted on a hand-packed trap column (Thermo Scientific POROS R2 reversed-phase resin 1112906, 1 mm ID × 2 cm, IDEX C-128) as described. Peptides were separated with a C18 analytical column (Waters 186002344), then eluted into the Q-Exactive Orbitrap Mass Spectrometer running in positive mode. On each experimental day, tandem mass spectrometry measurements were taken with undeuterated purified protein samples. Peptides were then identified using the Byonic software (Protein Metrics) with the HaloTag GT(GS)_3_ sequence as a reference sequence. Peptide lists were then exported and imported into HDExaminer 3 (Sierra Analytics) along with the deuterated and undeuterated sample mass spectra.

### HDExaminer 3 analysis, bimodal distribution fitting, and peptide classification

HDExaminer 3 was used to identify peptides in each HDX timepoint. A set of peptides that mapped primarily to one secondary structure was selected for detailed analysis, where peptides consistently showed low signal:noise ratios in the measured mass spectra. Peptide mass spectra were exported from HDExaminer 3, then unimodal and bimodal peaks were fit to Gaussian curves or to a sum of two Gaussians respectively as previously described (55). In refolding experiments, the centroid of each peak was allowed to drift to help identify peptides showing a burst in gain of protection between the unfolded control and the first deuteration timepoint.

In continuous labeling experiments with CP33 at 0M and 3.5M urea, peptides showing bimodal spectra in the 3.5M urea condition were considered structured in the CP33 intermediate. In experiments with CP217 ± TMR, peptides showing bimodal spectra or protection from exchange in the +TMR condition when compared to the −TMR condition were considered to be stabilized upon binding of TMR.

In pulsed-labeling experiments during protein refolding, peptides are considered to be fast-folding if they are >50% in the folded, less-deuterated peak at the 600s timepoint. This 600s time point was selected to coincide with the measured t_1/2_ of the fast-folding phase observed in HaloTag CD refolding experiments, and because an obvious difference was observed between fast- and slow-folding peptides at this time point. Peptides are considered part of the burst-phase intermediate if the unfolded peak shows a leftward shift of ≥ 0.5Da when comparing the unfolded control and the first 10s time point.

### Pulsed thiol-labeling during refolding

15μM protein was equilibrated overnight at 37 °C in 7.5M urea, in addition to a 1.6M urea HKMT refolding buffer and a 7.5M urea HKMT unfolded control buffer. To initiate refolding, protein was diluted twelve-fold into the refolding buffer to a concentration of 1.25μM, and at each refolding time point, protein was diluted further with a fluorescein-conjugated maleimide dye so 1μM protein was labeled with 50μM dye. The reaction was then quenched with 50mM DTT after 30s of labeling time. An unfolded control was prepared with the same experimental setup, using a 7.5M urea refolding buffer to dilute the unfolded protein sample. Timepoint samples and the unfolded control were mixed with SDS loading dye, boiled at 95C for 5 minutes, then run on a NuPAGE 4-12% Bis-Tris 1.5mm gel with an MES running buffer. Gels were then imaged for fluorescein fluorescence with an Alexa 488 filter set on a Typhoon gel scanner. In-gel fluorescence of protein bands were then quantified using ImageJ, and band intensities were normalized to the unfolded control. Data were then fitted with single-exponential or double-exponential kinetics as done previously (33) to measure rates of protection from labeling.

## Acknowledgements

We thank the Marqusee Lab for experimental advice and support, particularly HT Hobbs for advice with library generation and NGS and SM Costello and SR Shoemaker for assistance with HDX/MS protocols. We thank the Kuriyan Lab for assistance with FACS and NGS, VV Trotter (Deutschbauer Lab) for the Keio collection strains, and A Hung and KB Sander (Arkin Lab) for the ASKA expression vector and assistance with FACS. We thank F Ramirez and C Rose from QB3 Genomics for assistance with NGS. We thank W Coyote-Maestas assistance with structural parameter analyses. This work was supported by funding from the NIH (SM), the Chan Zuckerberg Biohub (SM), and the NSF Graduate Research Program Fellowship DGE 1752814 (NRD). SM is a Chan Zuckerberg Biohub Investigator.

## Author Contributions

NRD and SM designed research; NRD carried out all experimental studies, CATLM assisted with HDX/MS optimization, and HLTV performed initial CD experiments with CP33; NRD and SM analyzed data; NRD and SM wrote the paper.

