## Supplemental Figures and Tables for "The importance of the location of the N-terminus in successful protein folding *in vivo* and *in vitro*"

#### SI Fig. 1

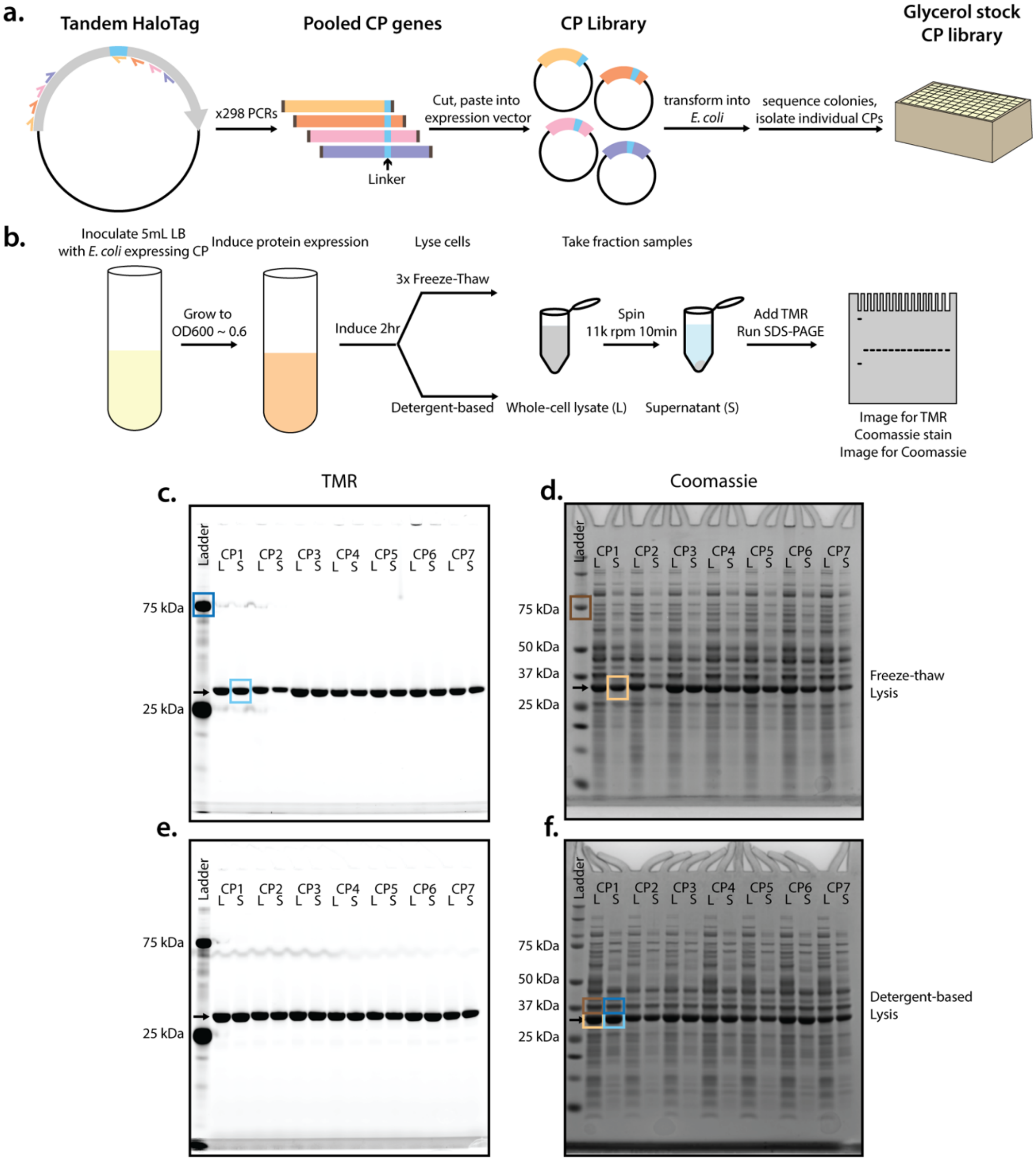


**Supplemental Figure 1. Overview of CP cloning and gel-based assay studying CP solubility and function.** a. Cloning schematic for generating all 297 CPs of HaloTag and isolating glycerol stocks of each CP. b. Methods schematic describing gel-based assay. A small culture of E. coli expressing a CP is grown to an OD600 of ~0.6, then induced with 1mM IPTG. Cells induce for another 2hr, then are lysed open with either three freeze-thaw cycles or with the detergent-based BugBuster protein extraction reagent. Samples of whole cell lysate (L) and the clarified supernatant (S) are taken and stained with the TMR ligand, then run via SDS-PAGE. Gels are imaged for TMR fluorescence, then stained with Coomassie. c-d. In-gel fluorescence and Coomassie staining of example gel for CPs 1-7 after freeze-thaw (FT) cell lysis. FT gel images were used to calculate functional scores for each CP, where the intensity of the protein band in the S fraction in the TMR image normalized by the intensity of a band in the ladder (blue squares) was divided by the intensity of protein in the S fraction in the Coomassie image normalized to the ladder (yellow, brown squares). e-f. In-gel fluorescence and Coomassie staining of example gel for CPs 1-7 after cell lysis via the detergent-based BugBuster reagent (DB). DB gel images were used to calculate solubility scores for each CP, where the intensity of the protein band in the S fraction normalized by the intensity of the band above it (blue squares) was divided by the intensity of protein in the L fraction normalized to the band above it (yellow, brown squares). Scores were then normalized to the CP1 score.

### SI Fig 2

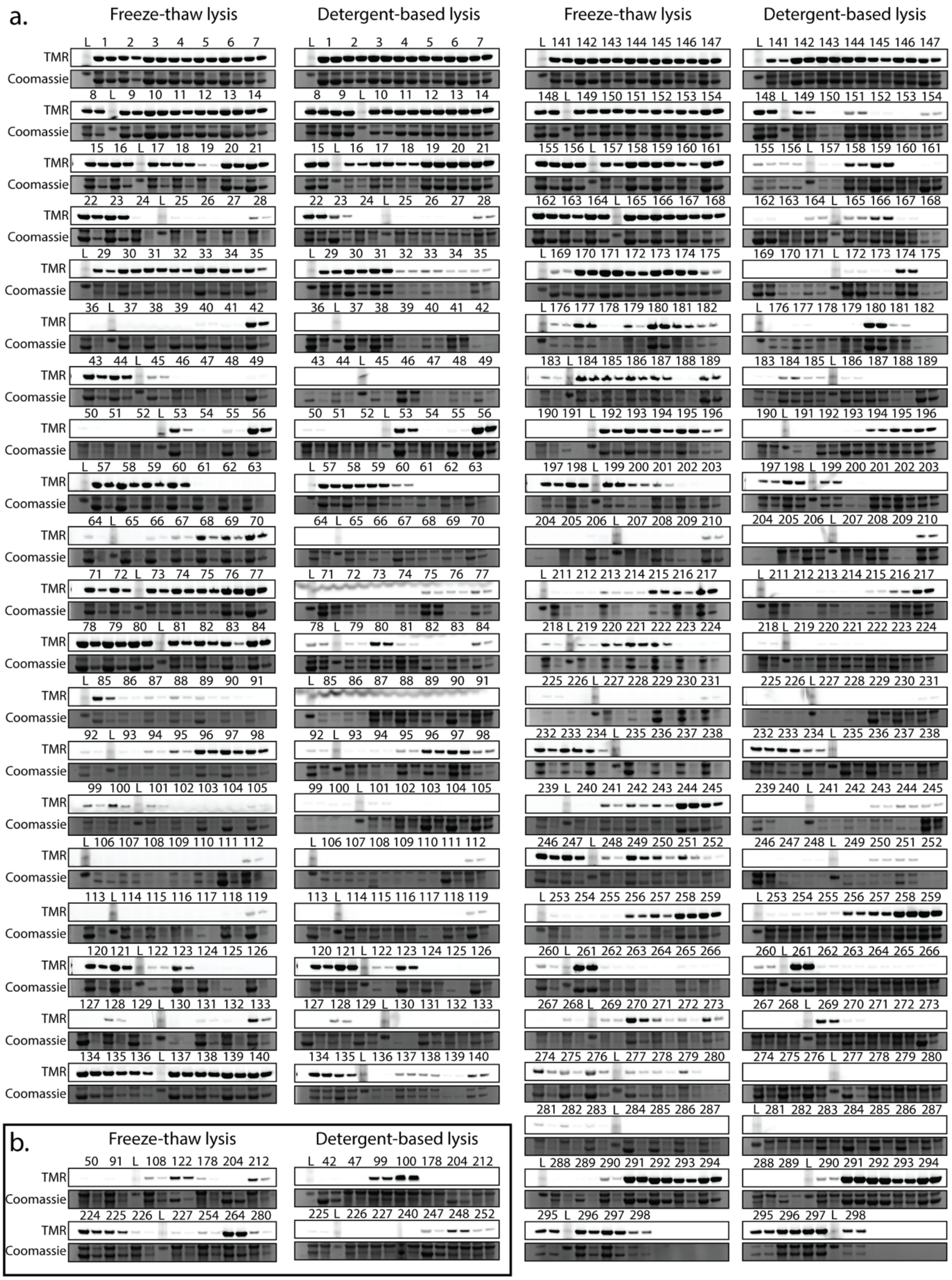
a.
Freeze-thaw lysis
Detergent-based lysis
Freeze-thaw lysis
Detergent-based lysis
TMR
Coomassie
b.
Freeze-thaw lysis
Detergent-based lysis
TMR
Coomassie

Supplemental Figure 2. Gel-based assay determining the solubility and function of every HaloTag circular permutant in *E. coli* cell lysate. **a.** Representative gels for every CP, using either the freeze-thaw or detergent-based lysis methods. For each CP, the left lane contains the whole cell lysate, while the right lane contains the soluble fraction as shown in Supp. Fig. 1c-f. “L”, ladder. “CP298” is WT HaloTag with the GS linker located at the N-terminus. **b.** Replicates of select CPs from gels in a. where cells grew poorly.

**SI Fig 3**

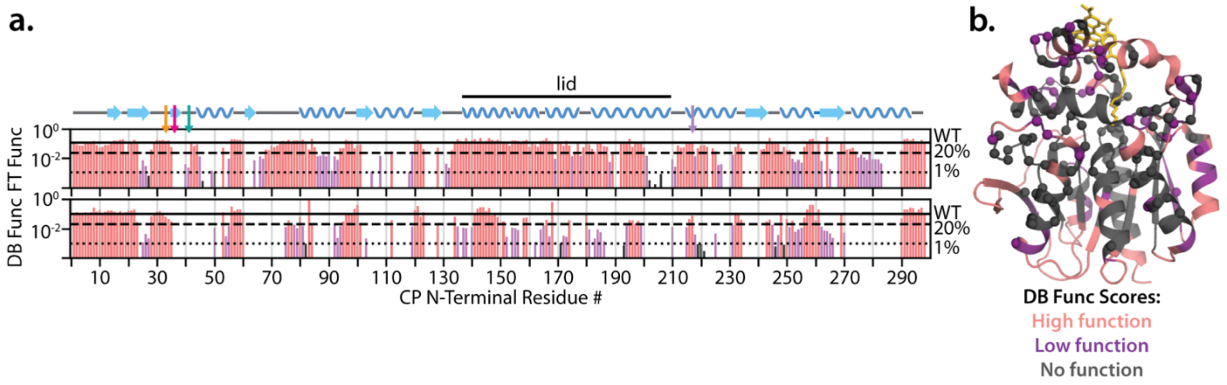


**Supplemental Figure 3. Quantification of CP function in gel-based assay using either the freeze-thaw or detergent-based lysis methods.** a. Functional scores from quantification of gel assays for every possible CP using a freeze-thaw lysis condition (FT) or a detergent-based lysis condition (DB). Blue dots above the DB Func data indicate CPs that are nonfunctional in BL21 FACS-seq data. One-dimensional topology map is shown above, with colored arrows indicating positions of CPs highlighted in Fig. 1b. Lines indicate the WT score(-), 20% of WT (--), and 1% of WT (••). High-function CPs have scores above the 20% line (pink), low-function CPs have scores between 1-20% the WT score (purple), and nonfunctioning CPs have scores of 0-1% the WT-level (black or no bar). b. CP locations with high (light pink), low (purple), and no protein function (dark gray) in DB lysis conditions. Spheres indicate the 118 CP positions with a decrease in function and change in categorization between FT and DB experiments.

### SI Fig 4

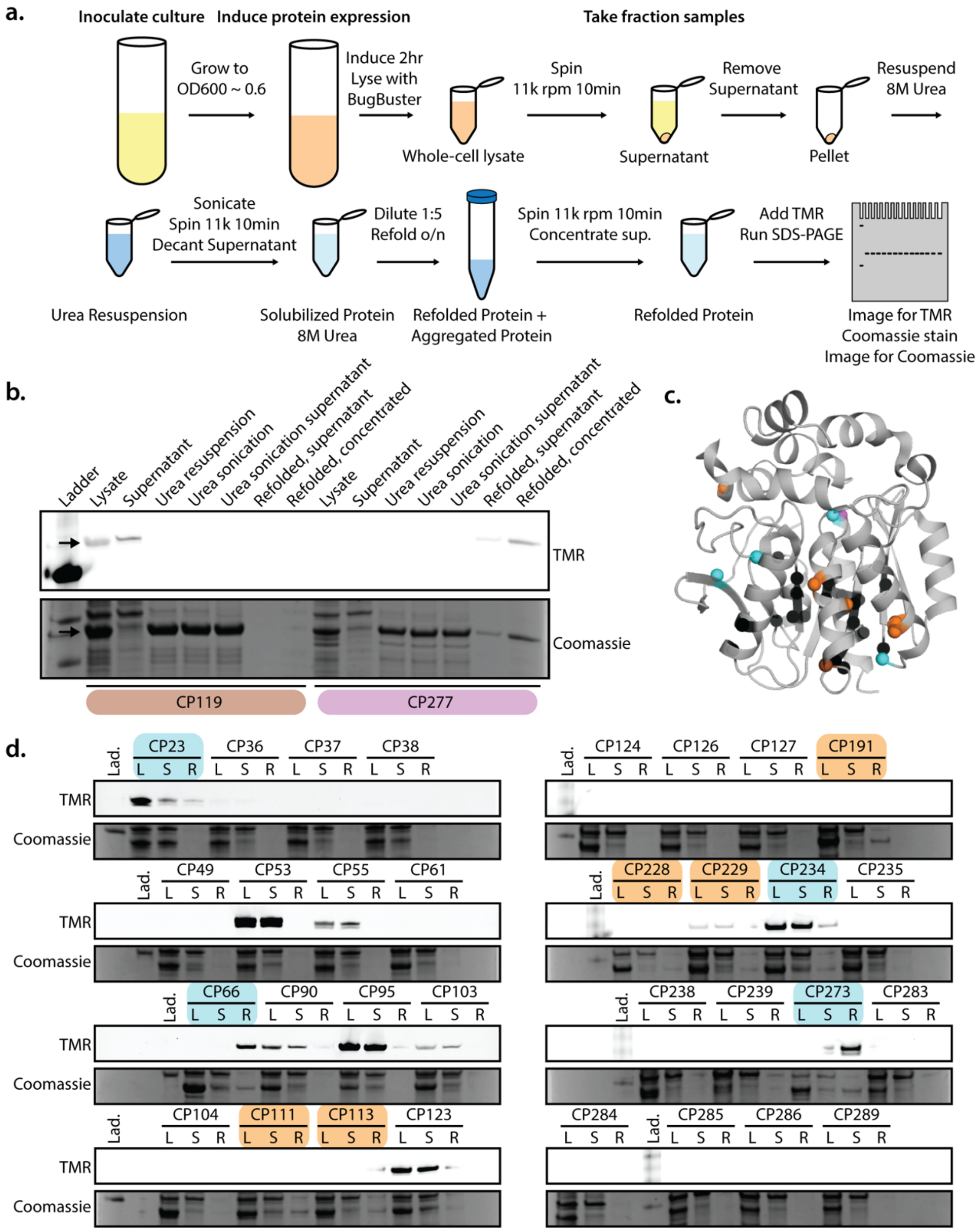


**Supplemental Figure 4. Testing the successful refolding of CP119 and CP277 from inclusion bodies.** a. Schematic outlining the inclusion body refolding assay. b. Gel-based assay examining whether CPs can be solubly refolded from inclusion bodies. CP119 does not refold at 1.6M urea, 37 °C and produce soluble, functional protein, while CP277 does. c. N-termini locations shown in spheres for CP119 (brown) and CP277 (purple). Refolded proteins in d that are in functional soluble (cyan), nonfunctional soluble (orange), and insoluble (black) categories. d. Additional CPs examined for successful refolding from inclusion bodies. CPs 23, 66, 234, and 273 refold solubly and can bind TMR (blue labels). CPs 111, 113, 191, 228, and 229 all yield a final soluble protein product, but protein may be misfolded or destabilized at 1.6M urea as no TMR binding is observed (orange labels). Lane labeling: L, whole cell lysate; S, supernatant; R, refolded and concentrated protein; Lad., ladder.

**SI Fig 5**

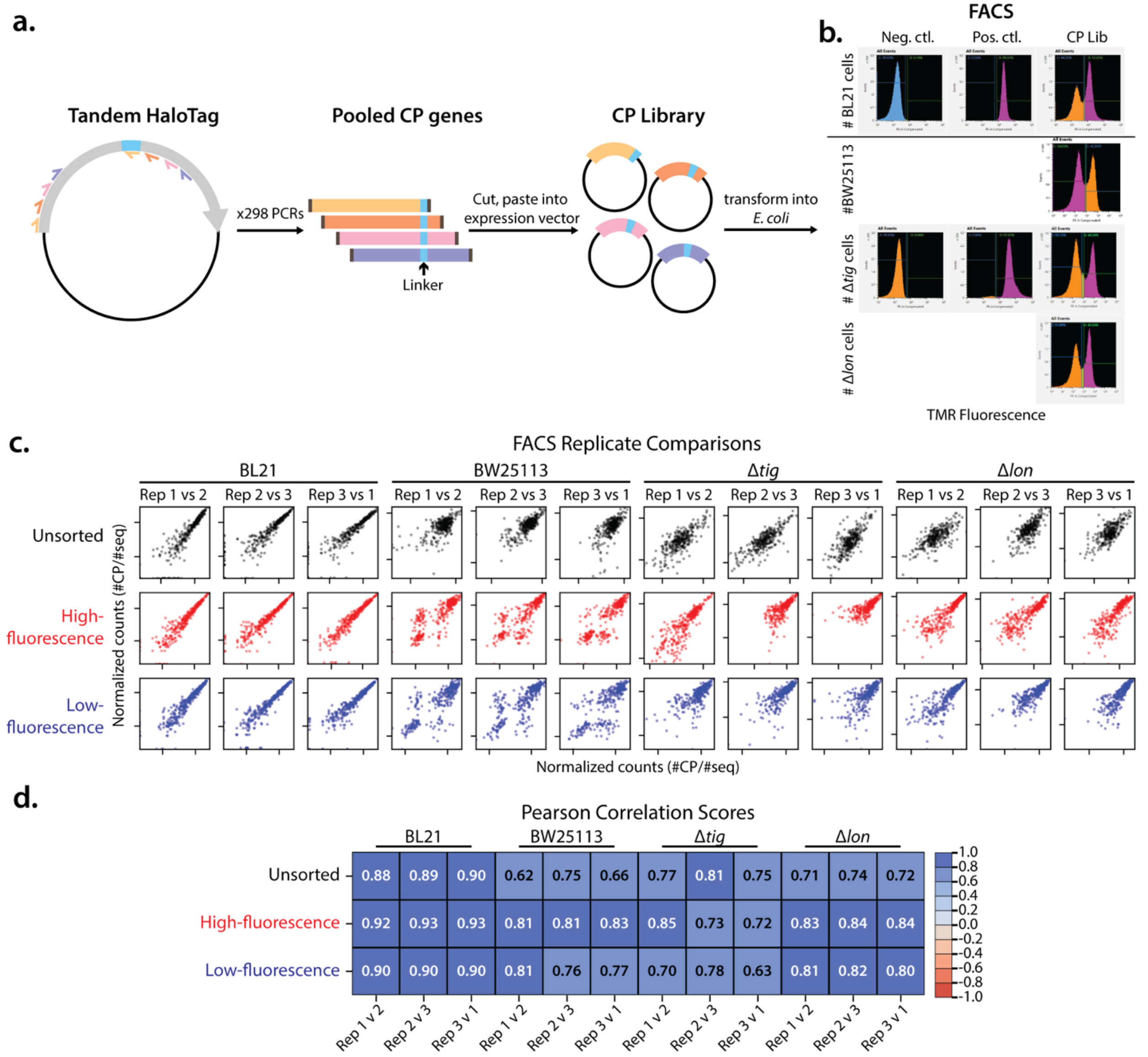


**Supplemental Figure 5. CP Library cloning strategy and representative FACS plots.** a. Pooled CP genes were cut and pasted into the BL21 expression vector or the ASKA expression vector for transformation into electrocompetent cells and preparation for sorting. b. FACS plots for BL21, BW25113, Δ*tig*, and Δ*lon* cells expressing the CP Library. The BL21 negative control consisted of cells expressing RNaseH, while the positive control was cells expressing HaloTag. For the three Keio strains, Δ*tig* cells expressing transformed with empty vector or cells expressing HaloTag were used for the negative and positive controls, respectively. c. Pairwise comparisons of normalized sequencing counts for each CP between replicates. “Unsorted” replicates are unsorted input sequences, “High-fluorescence” replicates are high TMR-fluorescence cell populations, while “Low-fluorescence” replicates are low TMR-fluorescence populations. d. Pearson correlation scores for the pairwise comparisons shown in c.

**SI Fig 6**

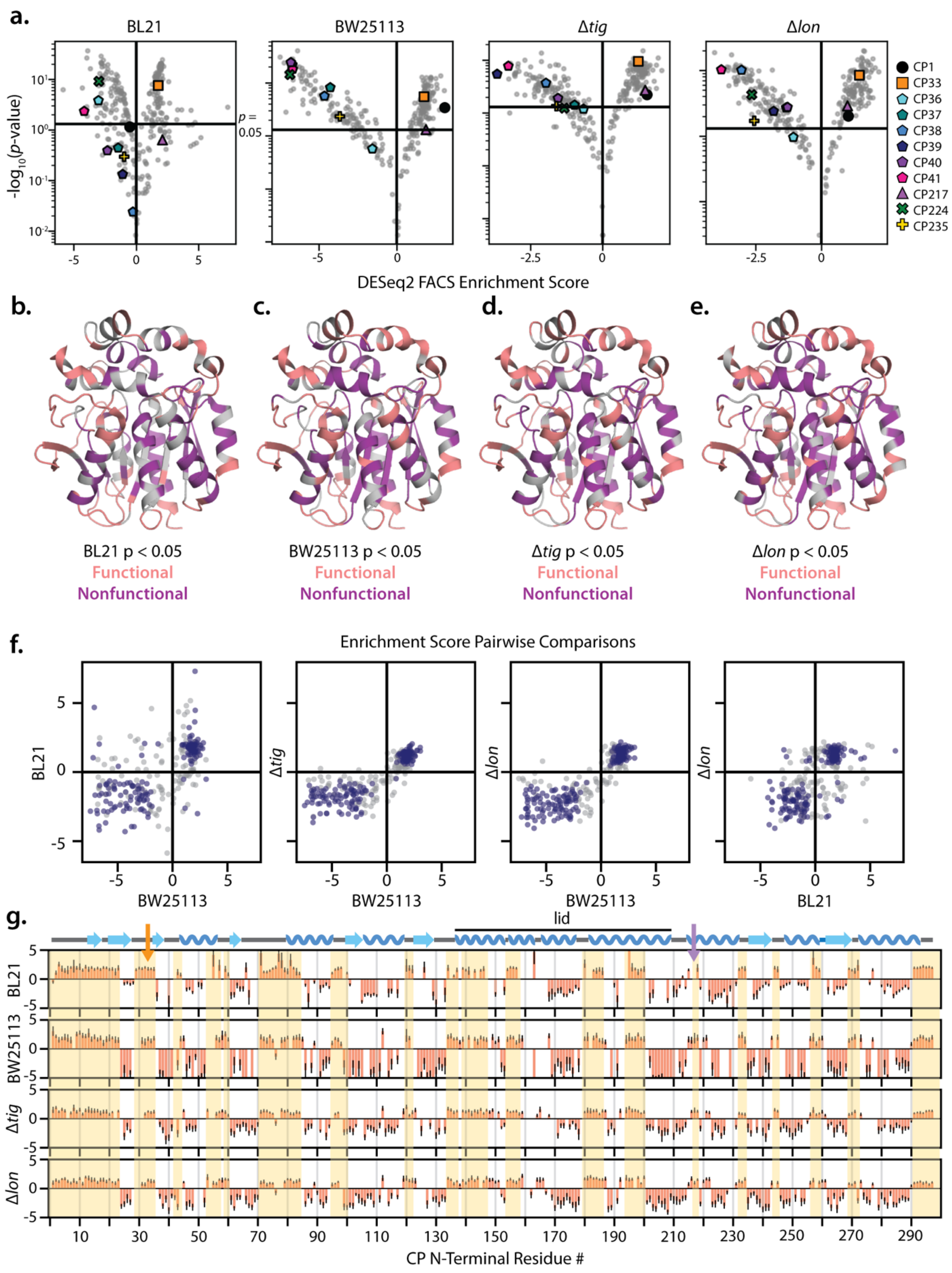
a.
BL21
BW25113
Δtig
Δlon
-log10(p-value)
p = 0.05
DESeq2 FACS Enrichment Score
CP1
CP33
CP36
CP37
CP38
CP39
CP40
CP41
CP217
CP224
CP235
b.
c.
d.
e.
BL21 p < 0.05
BW25113 p < 0.05
Δtig p < 0.05
Δlon p < 0.05
Functional
Nonfunctional
f.
Enrichment Score Pairwise Comparisons
BL21
Δtig
Δlon
BW25113
g.
lid
CP N-Terminal Residue #

**Supplemental Figure 6. DESeq2 statistical significance measurements for CP FACS enrichment scores.** a. Volcano plots for BL21, BW25113, Δtig, and Δlon FACS-seq enrichment scores. Horizontal lines represent $p < 0.05$ cutoff, and CPs with enrichment scores above this line are significantly enriched for function/nonfunction. Example CPs are also highlighted on each plot: CP1, C33, and CP217 are known to function in BL21 E. coli lysate gel assays, while CPs 36-41, 224, and 235 are all known to show low or no functional levels. b-e. CP positions with statistically-significant enrichment scores in each dataset. Functional CPs are shown in pink, while nonfunctional CPs are shown in purple. f. Pairwise comparisons between FACS-seq enrichment scores. Dark blue points represent CPs with significant enrichment scores in both datasets being compared, while grey points are nonsignificant in one or both datasets. CPs in the upper right and lower left quadrants are consistently functional or nonfunctional between datasets respectively. g. Bar graphs representing DESeq2 enrichment scores calculated for each CP where $p < 0.05$ in the high-fluorescence gate (error bars represent ± SE). Yellow boxes are the same as in Fig. 2a.

**SI Fig 7**

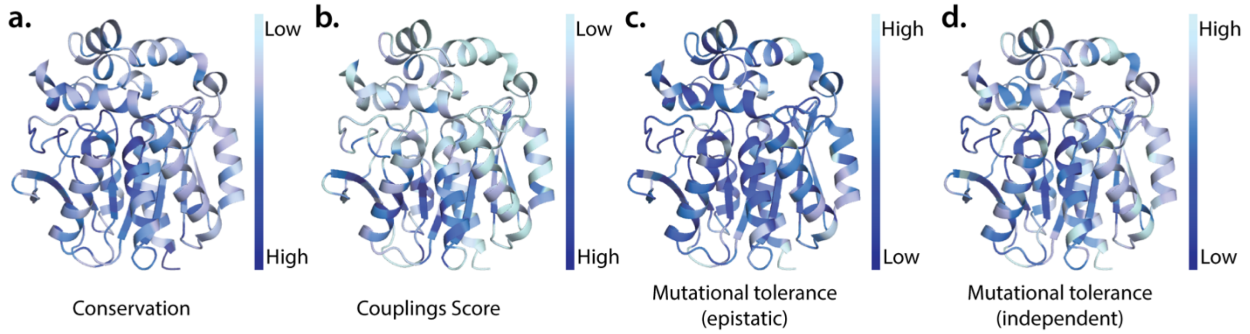


**Supplemental Figure 7. EVCouplings analyses of sequence conservation and mutational tolerance in HaloTag.** a. Per-residue sequence conservation plotted on the HaloTag structure (PDB: 6u32) based on an alignment of 4850 related protein sequences. b. EVCouplings couplings scores. c-d. Mutational tolerance of HaloTag calculated with the EVCouplings epistatic (c) and independent (d) models.

**SI Fig 8**

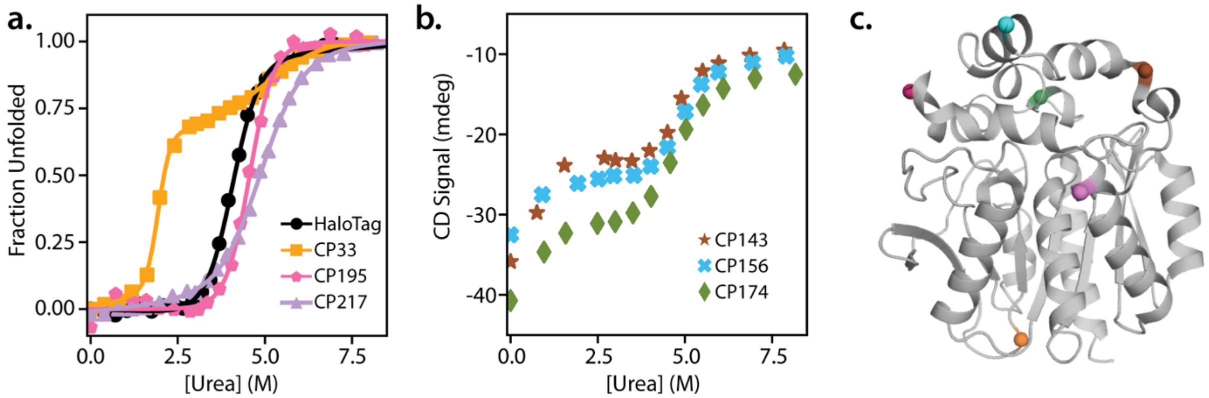


**Supplemental Figure 8. Urea denaturation of CPs as monitored by circular dichroism spectroscopy.** a. CP33 shows two unfolding transitions, while HaloTag, CP195, and CP217 only show one. CP195 is a lid CP with a similar *m*-value to WT HaloTag but a higher $C_m$. Fit values are in Supp. Table 1. b. CP143, 156, and 174 all have termini inserted in the lid, and contain what is likely a destabilized lid region that unfolds under low concentrations of urea. The second unfolding transition occurs with a Cm similar to that observed in CP33 and in CP217. c. N-termini locations of CPs 33, 143, 156, 174, 195, and 217 shown in spheres (orange, brown, blue, green, pink, and purple, respectively).

**SI Fig 9**

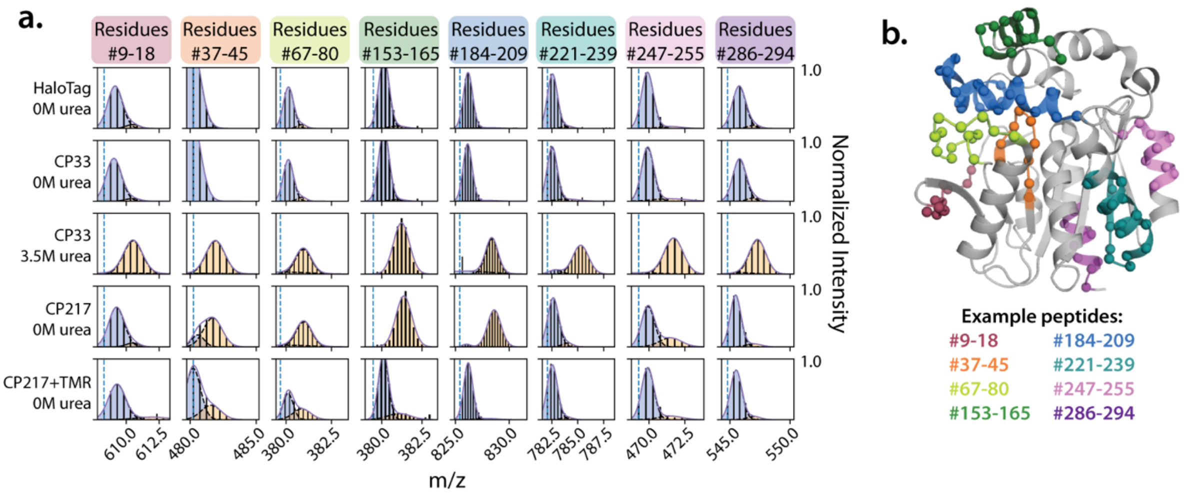


**Supplemental Figure 9. Continuous-labeling HDX/MS data for additional example peptides.** a. Peptides from purified HaloTag, CP33 in 0M and 3.5M urea, and CP217 with and without TMR after 10s of HDX. Peptides show high protection from exchange in HaloTag and CP33 in the absence of urea. When equilibrated at 3.5M urea, all of the example peptides in CP33 are fully-exchanged except for peptide 221-239, which shows a small population protected from exchange. Some core peptides (37-45 and 67-80) and all lid peptides (153-165 and 184-209) are fully exchanged in CP217, but are highly stabilized and show less exchange when TMR is bound. Other core peptides (247-255) show small amounts of stabilization when TMR is bound. b. Locations of example peptides mapped onto the HaloTag structure.

**SI Fig 10**

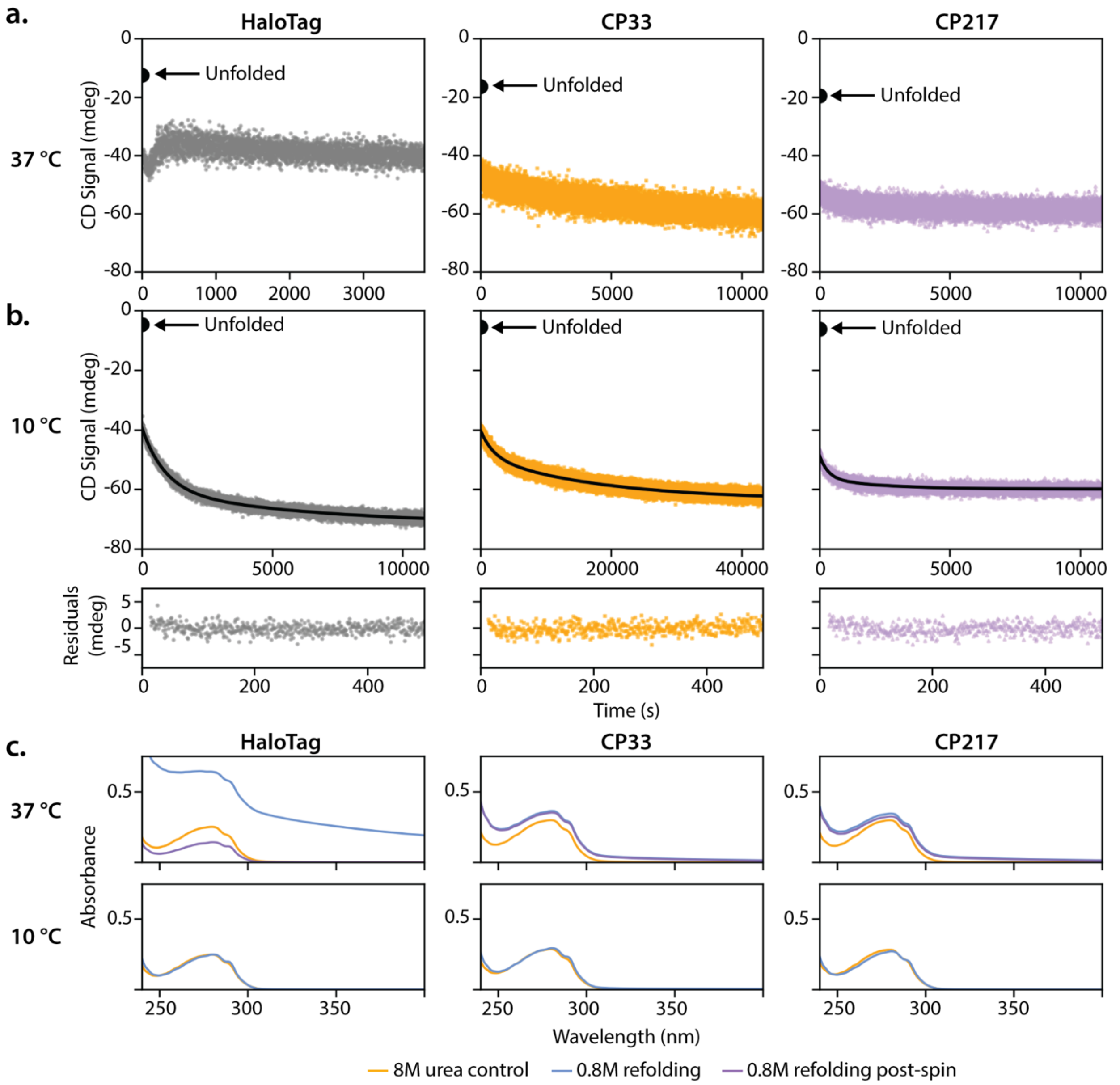


**Supplemental Figure 10. HaloTag, CP33, and CP217 are all aggregation-prone when refolding at 37 °C and at 0.8M urea.** a. Raw CD signal traces for 37 °C refolding experiments. HaloTag is more aggregation-prone than CP33 and CP217. After a burst phase decrease in CD signal, refolding HaloTag has a decrease in CD signal followed by an increase. This indicates aggregate formation. CP33 and CP217 only show a gradual decrease in CD signal. b. Raw CD signal traces, fits, and fit residuals for the 10 °C refolding experiments in Fig. 3.8. c. UV-Vis spectra of recovered CD samples. Spectra were taken of the unfolded proteins in 8M urea (orange lines) and the refolded proteins in 0.8M urea (blue lines) and buffer-corrected with the appropriate refolding buffers. If absorbance was observed between 300nm and 400nm in the 0.8M refolding samples, samples were spun at 16,000rpm for 10min. Another spectrum was taken of the supernatant (purple lines). At 37 °C, HaloTag shows high levels of absorbance between 300-350nm, and only

60% of the protein remained soluble after a hard spin. CP33 and CP217 show small amounts of absorbance between 300-350nm, but the aggregate did not pellet with the hard spin. At 10 °C, all three proteins refolded solubly.

### SI Fig 11

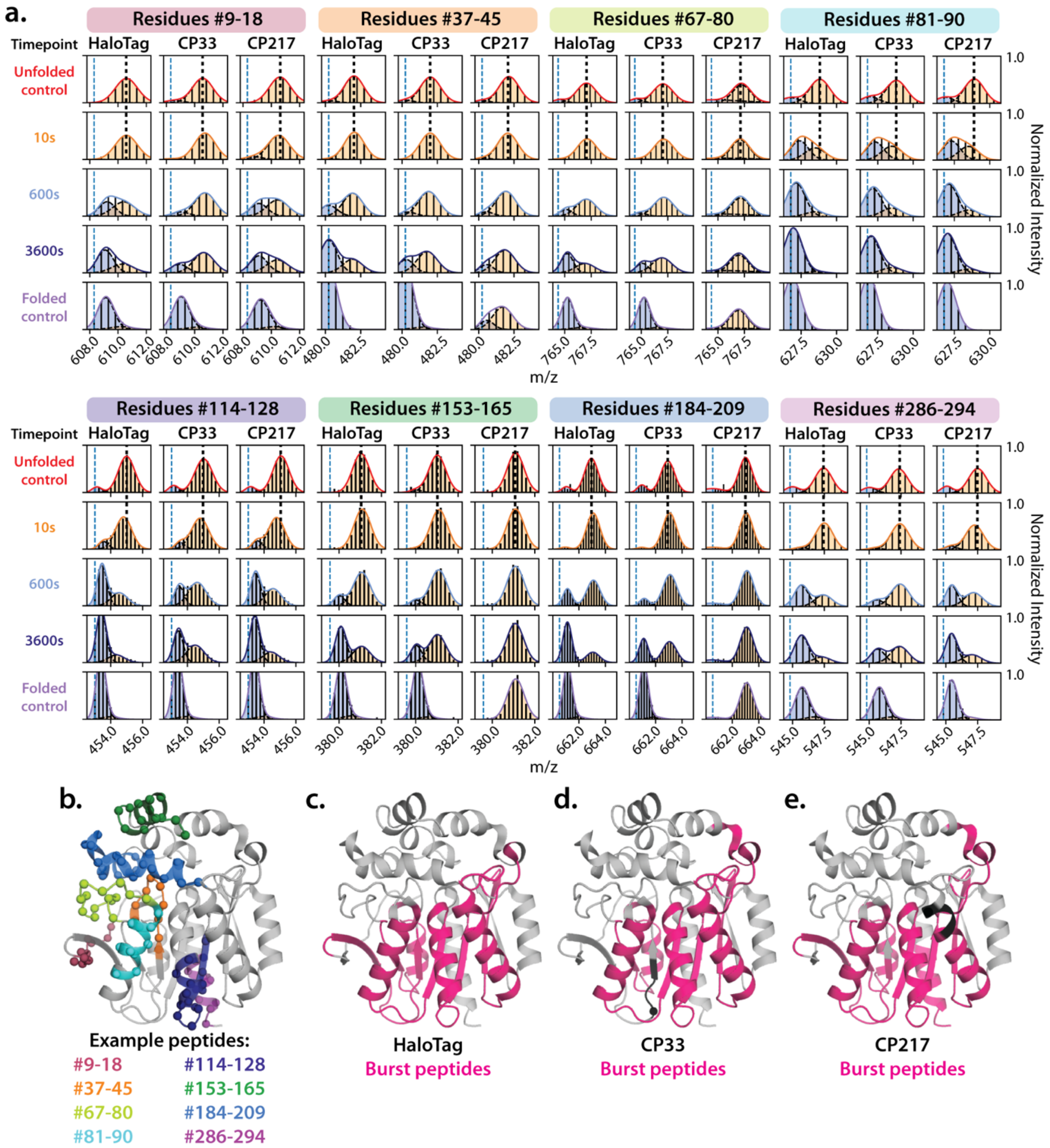


**Supplemental Figure 11. Example peptides from pulsed-labeling HDX/MS experiments during refolding and characterization of the burst phase intermediate.** a. Example refolding time courses for peptides from HaloTag, CP33, and CP217. Vertical black dotted lines are drawn based on the centroid of the peak in the unfolded control, and peptides where the unfolded peak in the 10s refolding time point shifts to the left (gaining protection) are considered part of the burst phase intermediate. The burst phase peptides shown here are 81-90 (1.5 Da) and 114-128 (1 Da), and in CP217, 286-294 (0.5 Da). b. Example peptides mapped onto the HaloTag structure. c-e. Burst phase peptides for HaloTag, CP33, and CP217. The burst phase is similar in each protein, except for residues 19-32 in CP33 (which is now a C-terminal peptide that no longer collapses in the burst phase), and residues 286-294 (which show a new burst-phase collapse that is not observed in HaloTag or in CP33). Peptides 46-66 and 221-239 are shown in Fig. 6a, and also show a burst phase collapse in all three proteins. 46-66 is then slow-folding in CP33 after the collapse.

SI Fig 12

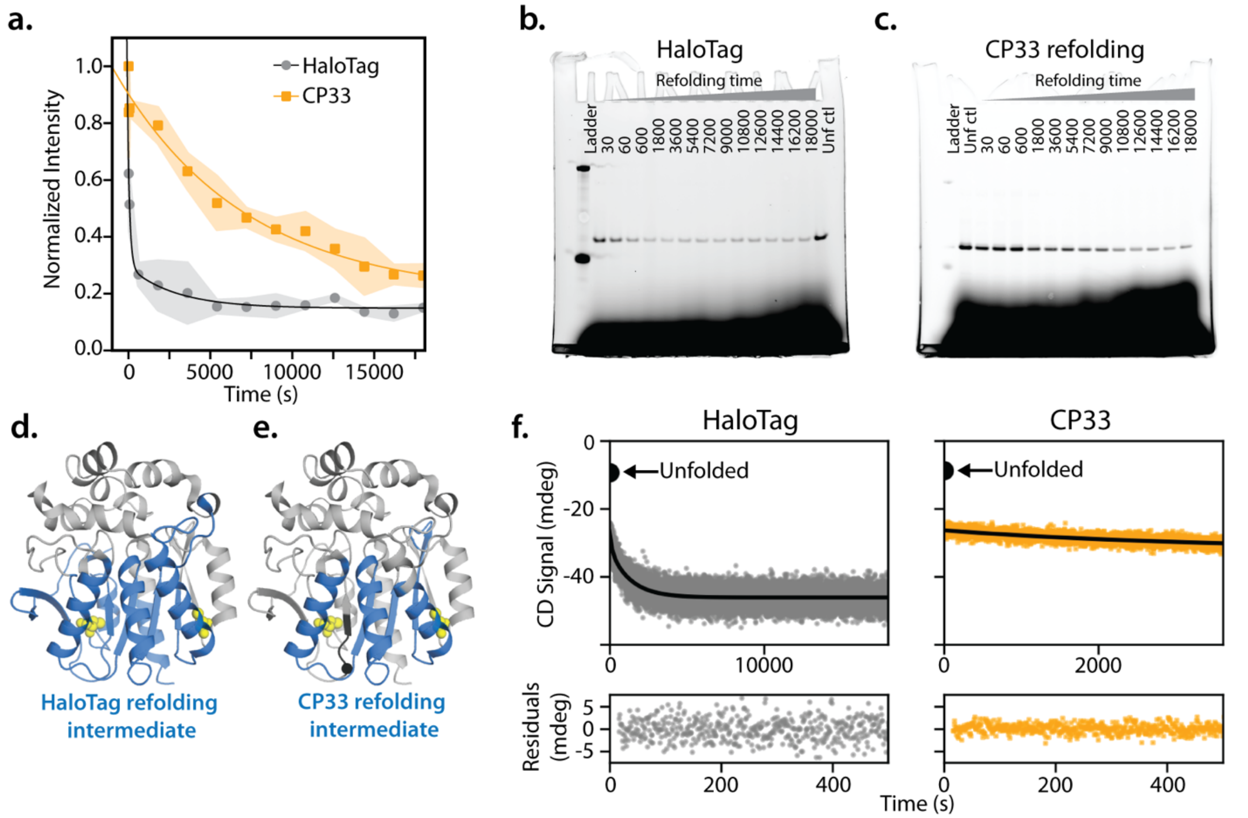


**Supplemental Figure 12. Pulsed-thiol labeling during refolding shows a change in refolding trajectory in CP33.** a-c. Quantification of gel-based refolding assay fit with biphasic kinetics (HaloTag, black line) or single-phase kinetics (CP33, orange line). Unfolded protein equilibrated at 7.5M at 37 °C was diluted to 1.6M urea to initiate refolding. At each timepoint, a sample was pulsed with a fluorescein-conjugated maleimide dye for 30s before labeling was quenched with 50mM DTT. Samples were run via SDS-PAGE and quantified for in-gel fluorescence (b-c), which was then normalized to the signal of an unfolded protein control. Shading represents standard deviation of three gel replicates, and b and c show two representative gels. d-e. Refolding intermediates identified by HDX/MS for HaloTag and CP33, with the native cysteines C61 and C262 represented by yellow spheres. f. Refolding at 37 °C and 1.6M urea monitored by CD at 225nm.

**SI Table 1**

| | HaloTag | CP33 | CP195 | CP217 |
|---|---|---|---|---|
| $\Delta G_{1,fold}$ (kcal/mol) | - | -5.71 ± 0.32 | - | - |
| $m_1$ (kcal/mol•M) | - | 2.99 ± 0.30 | - | - |
| $C_{m,1}$ (M) | - | 1.91 ± 0.02 | - | - |
| $\Delta G_{2,fold}$ (kcal/mol) | -6.42 ± 0.43 | -3.78 ± 0.60 | -7.04 ± 0.72 | -5.21 ± 0.22 |
| $m_2$ (kcal/mol•M) | 1.58 ± 0.11 | 0.76 ± 0.21 | 1.52 ± 0.16 | 1.06 ± 0.05 |
| $C_{m,2}$ (M) | 4.06 ± 0.54 | 4.97 ± 0.39 | 4.63 ± 0.88 | 4.92 ± 0.27 |

Supplemental Table 1. Fit parameters for urea denaturation of CPs shown in Supp. Fig. 8a. Errors report on one standard deviation of the fit parameters.

**SI Table 2**

| | HaloTag | CP33 |
|---|---|---|
| $t_{1/2,fast,CD,37C}$(s) | 105 ± 21 | - |
| $t_{1/2,slow,CD,37C}$ (s) | 917 ± 21 | 2264 ± 255 |
| $t_{1/2,fast,thiol,37C}$ (s) | 88 ± 23 | - |
| $t_{1/2,slow,thiol,37C}$ (s) | 2014 ± 717 | 5515 ± 1247 |

Supplemental Table 2. Refolding kinetics measured by CD and by pulsed-thiol labeling at 37 °C, 1.6M urea. Errors report on one standard deviation of the fit parameters.
